# Chaperone-mediated ordered assembly of the SAGA and NuA4 transcription co-activator complexes

**DOI:** 10.1101/524959

**Authors:** Alberto Elías-Villalobos, Damien Toullec, Céline Faux, Martial Séveno, Dominique Helmlinger

## Abstract

Transcription initiation involves the coordinated activities of large multimeric complexes that are organized into functional modules. Little is known about the mechanisms and pathways that govern their assembly from individual components. We report here several principles governing the assembly of the highly conserved SAGA and NuA4 co-activator complexes. Using fission yeast, which contain two functionally non-redundant paralogs of the shared Tra1 subunit, we demonstrate that Tra1 contributes to scaffolding the entire NuA4 complex. In contrast, within SAGA, Tra1 specifically promotes the incorporation of the de-ubiquitination module (DUB), defining an ordered assembly pathway. Biochemical and functional analyses elucidated the mechanism by which Tra1 assemble differentially into SAGA or NuA4 and identified a small, conserved region of Spt20 that is both necessary and sufficient to anchor Tra1 within SAGA. Finally, we establish that Hsp90 and its cochaperone TTT are required for Tra1 *de novo* incorporation into both SAGA and NuA4, indicating that Tra1, a pseudokinase of the PIKK family, shares a dedicated chaperone machinery with its cognate kinases. Overall, our work brings mechanistic insights into the *de novo* assembly of transcriptional complexes through ordered pathways and reveals the contribution of dedicated chaperones to this process.

## Introduction

A critical step in gene expression is transcription initiation, which is controlled by many factors that typically function as part of multimeric complexes. Genetic, biochemical, and structural evidence indicate that their subunits form distinct modules with specific functions and numerous studies have characterized their regulatory activities and roles in gene expression. In contrast, much less is known about how these complexes assemble, which chaperones are required, and whether their assembly can be modulated to control or expand their functions. Deciphering these principles is, however, important to understand their structural organization, function, and allosteric regulation (Marsh & Teichmann, 2015). Notably, chromatin-modifying and – remodeling complexes often share functional modules and therefore probably require dedicated mechanisms and chaperones for their proper assembly (Helmlinger & Tora, 2017).

One such complex, the Spt-Ada-Gcn5 acetyltransferase (SAGA) co-activator, bridges promoter-bound activators to the general transcription machinery. In yeast, SAGA is composed of 19 subunits, which are organized into five modules with distinct regulatory roles during transcription (Koutelou *et al*, 2010; Spedale *et al*, 2012). These include histone H3 acetylation (HAT), histone H2B de-ubiquitination (DUB), and regulation of TBP recruitment at core promoters. A fourth module consists of a set of core subunits that scaffold the entire complex, most of which are shared with the general transcription factor TFIID. Finally, its largest subunit, Tra1, directly binds to a diverse range of transcription factors. Tra1 is shared with another transcriptional co-activator complex, yeast NuA4, which also contains a HAT module that preferentially targets histone H4 and the H2A.Z variant (Lu *et al*, 2009).

Yeast Tra1 and its human ortholog, TRRAP, belong to a family of atypical kinases, the phosphoinositide 3 kinase-related kinases (PIKKs), but lack catalytic residues and therefore classify as pseudokinases (McMahon *et al*, 1998; Saleh *et al*, 1998; Vassilev *et al*, 1998). The reason for the evolutionary conservation of a typical PIKK domain architecture within Tra1 orthologs remains obscure. Genetic and biochemical studies indicate that Tra1 primary role is to recruit SAGA and NuA4 to specific promoters upon activator binding. It has been difficult, however, to delineate the specific contribution of Tra1 to SAGA and NuA4 architecture and activities because, to date, no clear separation-of-function alleles exist. How Tra1 interacts differentially with SAGA and NuA4 remains indeed poorly understood.

The fission yeast *Schizosaccharomyces pombe* provides a unique opportunity to address this issue because it has two paralogous proteins, Tra1 and Tra2, and each has non-redundant roles that are specific for SAGA and NuA4, respectively (Helmlinger, 2012). Within SAGA, Tra1 has specific regulatory roles and does not contribute to its overall assembly (Helmlinger *et al*, 2011), consistent with its peripheral position in the recent cryo-electron microscopy structure of SAGA from the budding yeast *Pichia pastoris* (Sharov *et al*, 2017). In contrast, a recent partial structure of the yeast NuA4 complex indicates that Tra1 occupies a more central position (Wang *et al*, 2018). However, little is known about how Tra1 incorporates into the SAGA and NuA4 complexes, whether it involves similar or distinct mechanisms, and which chaperone or assembly factors are required.

Here, we addressed these issues and show that, in *S. pombe*, Tra1 and Tra2 require Hsp90 and its cochaperone, the Triple-T complex (TTT), for their *de novo* incorporation into the SAGA and NuA4 complexes, respectively. Furthermore, proteomic, biochemical, and genetic approaches identified the residues that mediate Tra1 specific interaction with SAGA, which contacts a remarkably small region of the core subunit Spt20. Kinetic analyses of nascent Tra1 incorporation revealed that it promotes the incorporation of the DUB module into SAGA, uncovering an ordered pathway of SAGA assembly. Finally, in contrast to Tra1 within SAGA, we show that Tra2 has a general scaffolding role in NuA4 assembly. Overall, our work brings mechanistic insights into the assembly and modular organization of two important transcriptional co-activator complexes.

## Results

Previous work in mammalian cells revealed that the Hsp90 cochaperone TTT stabilizes PIKKs, including TRRAP, the human ortholog of yeast Tra1 (Takai *et al*, 2007; Anderson *et al*, 2008; Takai *et al*, 2010; Hurov *et al*, 2010; Kaizuka *et al*, 2010; Izumi *et al*, 2012). Three specific subunits, Tel2, Tti1, and Tti2, define the TTT complex in *S. pombe, S. cerevisiae*, and human cells (Table S1) (Hayashi *et al*, 2007; Shevchenko *et al*, 2008; Takai *et al*, 2010). Tra1 interacts physically and genetically with some TTT subunits in *S. cerevisiae* and *S. pombe* (Hayashi *et al*, 2007; Shevchenko *et al*, 2008; Helmlinger *et al*, 2011; Inoue *et al*, 2017; Genereaux *et al*, 2012). Fission yeasts have two paralogous genes, *tra1*+ and *tra2*+, and each has non-redundant roles that are specific for SAGA or NuA4, respectively (Helmlinger *et al*, 2011). *S. pombe* thus offers a unique opportunity to study the specific roles of TTT and Tra1 in SAGA and NuA4 complex architecture and regulatory functions.

### The TTT subunit Tti2 contributes to Tra1 and Tra2 function in gene expression

We first determined the contribution of TTT to Tra1-SAGA and Tra2-NuA4 dependent gene expression in *S. pombe*. In human cells, TTI2 is critical for the stability of both TEL2 and TTI1 at steady state (Hurov *et al*, 2010), in agreement with its stable association within the TTT complex. We thus focused our analysis on Tti2, which we confirm interacts with both Tra1 and Tra2 (Table S1). In *S. pombe*, Tra2 is essential for viability, similar to Tti2, whereas *tra1*Δ mutants are viable (Helmlinger *et al*, 2011; Inoue *et al*, 2017). Using a strategy based on inducible CreER-loxP-mediated recombination, we generated conditional knock-out alleles of *tti2*+ (*tti2-CKO*) (Figure S1) and *tra2*+ (*tra2-CKO*) (Figure S2). Both strains showed β-estradiol-induced loss of Tti2 or Tra2 expression, accompanied by progressive proliferation defects.

We then performed genome-wide expression analyses of DMSO- and β-estradiol-treated *tti2-CKO* and *tra2-CKO* cells, compared to a *cre-ER* strain treated identically. We also analyzed *tra1*Δ mutants that were compared to a wild-type strain. Differential expression analysis revealed specific and overlapping changes in each mutant (Figure S3A-D). For example, we confirmed down-regulation of the *SCC569.05c*^+^ and *gst2*^+^ genes in both *tti2-CKO* and *tra1*Δ mutants (Figure S3E) and of *SPCC1884.01*^+^ and *SPCAC977.12*^+^ in both *tti2-CKO* and *tra2-CKO* mutants (Figure S3F). To compare their overall transcriptome profiles, we performed hierarchical clustering of all differentially expressed genes and found all possible classes. These include genes which expression is up- or down-regulated in only one mutant, in two mutants, or in all three mutants (Figure 1A). A Venn diagram representation of all differentially expressed genes showed the extent of the overlap between all three mutants. Remarkably, 105 out of the 184 Tti2-dependent genes are also regulated by Tra1 (25 genes), Tra2 (72 genes), or both Tra1 and Tra2 (8 genes) (Figure 1B). Finally, Tra1 has important roles in recruiting SAGA and NuA4 to chromatin. We thus evaluated the effect of Tti2 on the binding of the SAGA subunit Spt7 and the NuA4 subunit Epl1 to specific promoters, using chromatin immunoprecipitation (ChIP). Upon depletion of Tti2, we observed reduced occupancy of Spt7 at the *pho84*+ and *mei2+* promoters and of Epl1 at the *ssa2*+ promoter, despite normal steady-state levels (Figure 1C,D).

**Figure 1.**
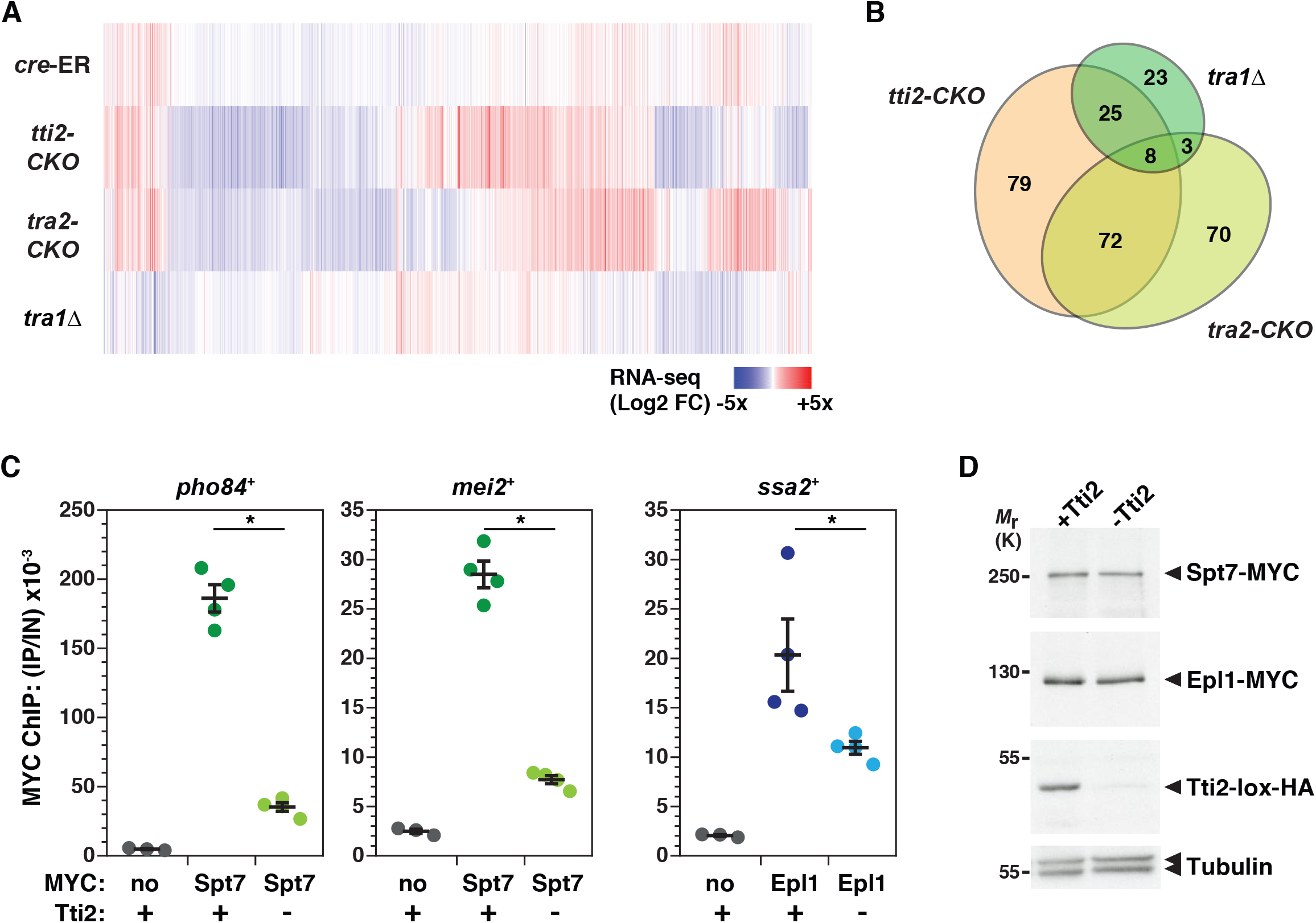
The Tti2 subunit of the TTT complex contributes to Tra1 and Tra2 functions in gene expression. (A) Hierarchical clustering analysis of the transcriptome profiles of control *cre-ER* (*cre* (WT)), inducible *tti2*+ (*tti2-CKO*) and *tra2*+ knock-outs (*tra2-CKO*), and *tra1*Δ mutants (rows). Differential gene expression analysis was performed comparing cells treated with either DMSO (control) or β-estradiol (KO) for 21 hours (h) (*cre-ER* and *tra2-CKO* strains) or 18 h (*tti2-CKO* strain). *tra1*Δ mutants were compared to isogenic wild-type (WT) control cells. Columns represent genes that are differentially expressed in at least one condition (*P* ≤ 0.01) and clustered based on Pearson distance. The Log2 of their fold-change in each condition is colour coded, as indicated. (B) Venn diagrams showing the overlap between genes that are differentially expressed (FC ≥ 1.5, *P* ≤ 0.01) in inducible *tti2-CKO* (n = 184), *tra2-CKO* (n = 153), and *tra1*Δ mutants (n = 59). Genes which expression is β-estradiol-regulated (see Figure S3A) have been filtered out before constructing the diagrams. (C) SAGA and NuA4 promoter binding upon inducible *tti2* knock-out. Chromatin immunoprecipitation followed by quantitative PCR (ChIP-qPCR) analysis was performed using *tti2-CKO* cells treated with either DMSO (+) or β-estradiol (-) for 18 h. ChIP of Spt7-MYC at the *pho84*+ and *mei2*+ promoters or of Epl1-MYC at the *ssa2*+ promoter serve as proxies for SAGA or NuA4 binding, respectively. A non-tagged strain was used as control for background IP signal (MYC: no). Ratios of MYC ChIP to input (IP/IN) from three independent experiments are shown as individual points, overlaid with the mean and SEM. Statistical significance was determined by one-way ANOVA followed by Tukey’s multiple comparison tests (\**P* < 0.05). (D) Anti-MYC and – HA Western blotting of Tti2-HA, Spt7-MYC, and Epl1-MYC in the input fraction of the chromatin samples used for the ChIP-qPCR experiments shown in (E). Equal loading was controlled using an anti-tubulin antibody.

In conclusion, we accumulated functional evidence suggesting that Tti2, likely as part of the TTT complex, contributes to the regulatory activities of Tra1 and Tra2 in gene expression. Therefore, similar to their active counterparts, the Tra1 and Tra2 pseudokinases require the TTT cochaperone to function.

### Tti2 promotes the *de novo* incorporation of Tra1 and Tra2 into SAGA and NuA4

These observations prompted us to test whether Tti2, as an Hsp90 cochaperone, promotes the incorporation of Tra1 and Tra2 into SAGA and NuA4, respectively, as shown for human mTOR and ATR-containing complexes (Takai *et al*, 2010; Kaizuka *et al*, 2010). For this, we affinity purified SAGA and NuA4 upon conditional deletion of *tti2*+. Silver staining and quantitative MS analyses revealed a 10-fold reduction of Tra1 from SAGA when Tti2 is depleted, as compared to control conditions (Figure 2A). Similarly, we observed about a 2-fold reduction in the levels of Tra2 in purified NuA4 complexes (Figure 2B).

**Figure 2:**
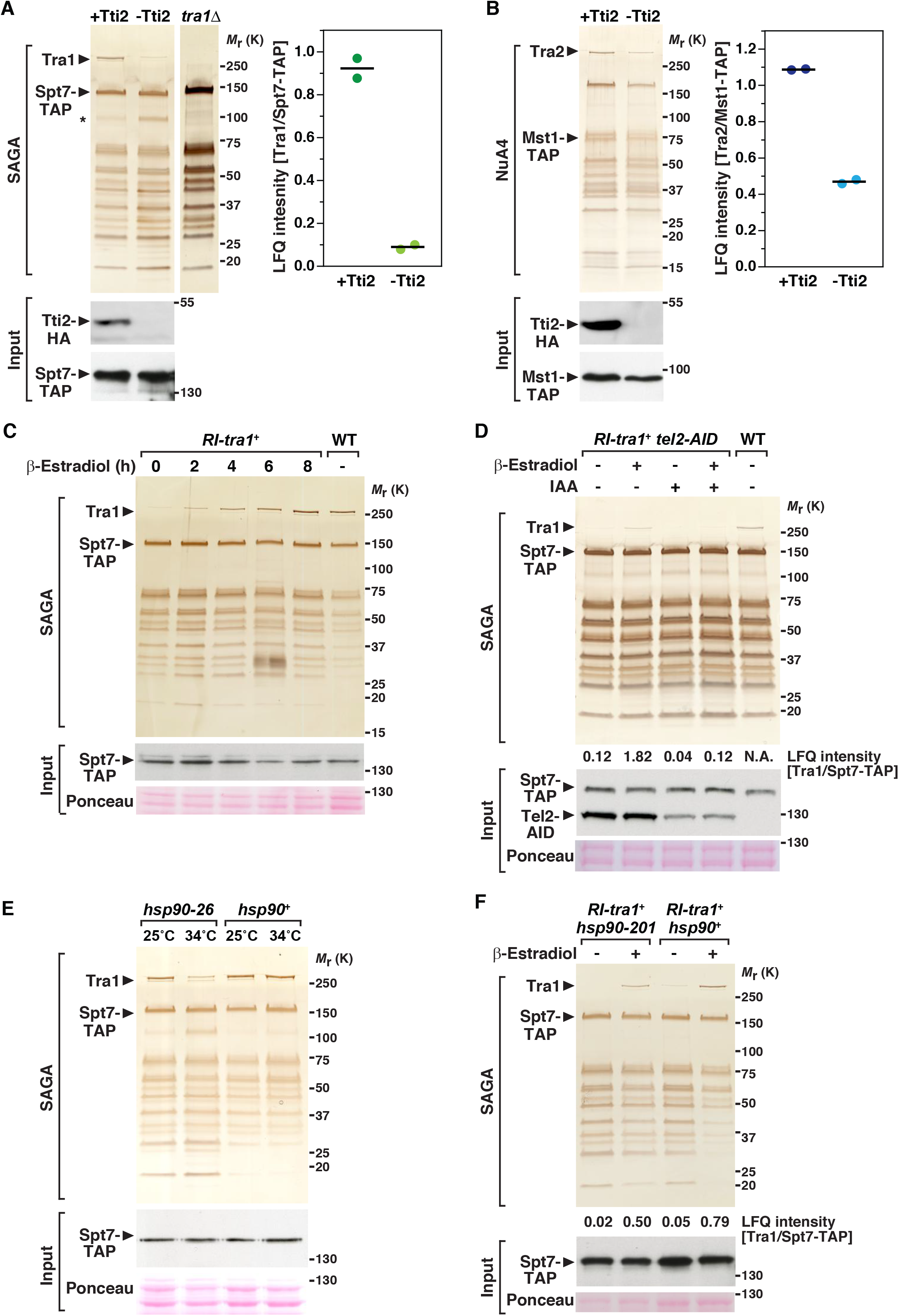
Hsp90 and Tti2 promote the incorporation of Tra1/Tra2 into SAGA/NuA4 complexes. (A) Silver staining of SAGA complexes purified either in the presence or absence of Tti2 (left). *spt7-TAP tti2-CKO* cells were grown to exponential phase in rich medium supplemented with either DMSO (+Tti2) or β-estradiol (-Tti2) for 18 h. SAGA was purified from a *tra1*Δ strain as a control for the complete loss of Tra1 from SAGA. LC-MS/MS analyses of SAGA purifications (right). LFQ intensity ratios of Tra1 to the bait, Spt7, from two biological replicates are plotted individually with the mean (black bar). Below are anti-HA Western blotting of Spt7-TAP and Tti2-HA in a fraction of the input used for TAP. (B) Silver staining of NuA4 complexes purified either in presence or absence of Tti2 (left). *mst1-TAP tti2-CKO* cells were grown to exponential phase in rich medium supplemented with either DMSO (+Tti2) or β-estradiol (-Tti2) for 18 h. LC-MS/MS analyses of NuA4 purifications (right). LFQ intensity ratios of Tra2 to the bait, Mst1, from two biological replicates are plotted individually with the mean (black bar). Below are anti-HA Western blotting of Mst1-TAP and Tti2-HA in a fraction of the input used for TAP. (C) Silver staining analysis of SAGA complexes purified upon Tra1 de novo expression (left). *spt7-TAP RI-tra1* cells were grown to exponential phase and harvested at different time-points after β-estradiol addition, as indicated (hours). SAGA was purified from a WT strain as a positive control. Below are anti-HA Western blot analyses of Spt7-TAP in a fraction of the input used for TAP. Ponceau red staining is used as loading control. Shown are data that are representative of four independent experiments. (D) Silver staining of SAGA complexes purified upon Tra1 *de novo* expression, either in presence or absence of Tel2. *spt7-TAP RI-tra1 tel2-AID* cells were grown to exponential phase in rich medium, supplemented with either ethanol (-IAA) or auxin (+IAA) for 16 h, and harvested 6 hours after addition of either DMSO (-) or β-estradiol (+). SAGA was purified from an untreated WT strain as a positive control. Numbers at the bottom of the gel represent LFQ intensity ratios of Tra1 to the bait, Spt7, from LC-MS/MS analyses of purified SAGA complexes (N.A.: not analyzed). Below are anti-HA Western blotting of Spt7-TAP and Tel2-AID in a fraction of the input used for TAP. Both the TAP and AID sequences are in frame with HA epitopes. Ponceau red staining is used as loading control. Shown are data that are representative of two independent experiments. (E) Silver staining analysis of SAGA complexes purified using Spt7 as the bait from WT and *hsp90-26* mutants. Cells were grown to exponential phase at the permissive temperature (25°C) and shifted for 6 hours at the restrictive temperature (34°C). Below are anti-HA Western blot analyses of Spt7-TAP in a fraction of the input used for TAP. Ponceau red staining is used as loading control. Shown are data that are representative of two independent experiments. (F) Silver staining of SAGA complexes purified upon Tra1 *de novo* expression, in WT and *hsp90-201* mutant strains. *spt7-TAP RI-tra1 hsp90*+ and *spt7-TAP RI-tra1 hsp90-201* cells were grown to exponential phase in rich medium and harvested 6 hours after addition of either DMSO (-) or β-estradiol (+). Numbers at the bottom of the gel represent LFQ intensity ratios of Tra1 to the bait, Spt7, from LC-MS/MS analyses of purified SAGA complexes. Below are anti-HA Western blotting of Spt7-TAP in a fraction of the input used for TAP. Ponceau red staining is used as loading control. Shown are data that are representative of two independent experiments.

We next tested if Tti2 prevents Tra1 and Tra2 disassembly from their complex or, rather, promotes their *de novo* incorporation. For this, we took advantage of the viability of *tra1*Δ mutants and disrupted the *tra1*+ promoter with a transcription terminator sequence flanked by *loxP* sites (*RI-tra1*+, Figure S4). With this allele, CreER-mediated recombination allows the inducible expression of Tra1 at endogenous levels. As a proof of principle, β-estradiol addition to *RI-tra1*+ strains restored their growth defects in conditions of replicative stress, using hydroxyurea (HU) (Figure S4), to which *tra1*Δ mutants are sensitive (Helmlinger *et al*, 2011). Purification of SAGA from *RI-tra1+* cells showed a time-dependent, progressive increase of Tra1 in Spt7 purification eluates upon β-estradiol addition, validating this approach for monitoring the *de novo* incorporation of Tra1 into SAGA (Figure 2C). However, to conditionally deplete TTT in *RI-tra1+* cells, we had to develop a strategy different from the CreER-loxP-mediated knockout used so far. Fusing the TTT subunit Tel2 to an auxin-inducible degron (AID) allowed its inducible degradation and caused subsequent proliferation defects by adding the plant hormone auxin (Figure S5A-C). Silver staining and quantitative MS analyses of SAGA purified from *RI-tra1*+ *tel2-AID* cells showed reduced interaction between *de novo* produced Tra1 and affinity purified Spt7 in cells partially depleted of Tel2 (lane 2 *vs* 4, Figure 2D). These results demonstrate that TTT contributes to the *de novo* incorporation of Tra1 into the SAGA complex.

Work in human cells revealed that TTT functions as an adaptor, which recruits the HSP90 chaperone to PIKKs specifically (Takai *et al*, 2010; Izumi *et al*, 2012; Pal *et al*, 2014). We thus determined if the *de novo* incorporation of Tra1 into SAGA requires Hsp90 in *S. pombe*. We first tested the effect of the conditional inactivation of Hsp90 on SAGA subunit composition at steady state. For this, we affinity purified Spt7 from *hsp90-26* temperature-sensitive mutants grown at either permissive or restrictive temperature (Aligue *et al*, 1994). Silver staining analysis showed that Hsp90 inactivation caused a specific decrease of Tra1 in Spt7 purification eluates (Figure 2E). We next asked whether Hsp90 contributes to *de novo* incorporation of Tra1 into SAGA. However, CreER cytoplasmic sequestration depends on a functional Hsp90. To minimize its impact on CreER, we used *hsp90-201* mutants, which harbor a weaker Hsp90 mutant allele (Alaamery & Hoffman, 2008). Silver staining and quantitative MS analyses revealed a decrease of newly synthesized Tra1 in SAGA purified from *hsp90-201* mutants, as compared to wild-type cells (lane 4 *vs* 2, Figure 2F). Although the observed effect is modest in this experimental condition, this result supports the conclusion that Hsp90, like TTT, contributes to *de novo* incorporation of Tra1 into SAGA. Altogether, these data indicate that TTT acts as an Hsp90 cochaperone that promotes the assembly of Tra1 into SAGA and likely Tra2 into NuA4. Therefore, pseudokinases and kinases of the PIKK family share a specific, dedicated chaperone machinery for their maturation and incorporation into active complexes.

### Tra1 and Tra2 have distinct architectural roles between SAGA and NuA4

We noted that the absence of Tti2 affected SAGA and NuA4 differently. Upon Tti2 depletion, the decrease of Tra1 does not affect SAGA overall migration profile, similar to what we observed in a *tra1*Δ mutant (Figure 2A) (Helmlinger *et al*, 2011). In contrast, the effect of Tti2 on Tra2 incorporation within NuA4 is less pronounced, but seems to cause a global decrease in the amount of purified NuA4 (Figure 2B). Alternatively, the bait used for this purification, the Mst1 HAT subunit, might dissociate from the rest of the complex upon *tti2*+ deletion and loss of Tra2.

To directly evaluate the effect of Tra2 on NuA4 subunit composition, we purified NuA4 upon *tra2*+ deletion, using *tra2-CKO* cells. For this, we affinity purified either Epl1, which anchors the HAT module to the rest of NuA4 in *S. cerevisiae* (Boudreault *et al*, 2003), or Vid21, which *S. cerevisiae* ortholog Eaf1 is a platform for NuA4 assembly (Auger *et al*, 2008). Silver staining and quantitative MS analyses of affinity purified Mst1, Epl1, and Vid21 revealed that each interact with a similar set of 13 proteins that define the NuA4 complex from *S. pombe* (Figure S6A-C), confirming and extending results from a previous study (Shevchenko *et al*, 2008). Upon the loss of Tra2, we observed an overall decrease in the amount of purified NuA4, using either Epl1 or Vid21 as baits (Figure 3A). Quantitative MS analyses of Vid21 purification eluates confirmed an overall decrease of all 13 NuA4 subunits upon the loss of Tra2 (Figure 3B). Altogether, our biochemical analyses indicate that, in contrast to Tra1 in SAGA, Tra2 contributes to the scaffolding and stabilization of the entire NuA4 complex. These distinct structural roles are consistent with the peripheral position of Tra1 within SAGA and the central position in NuA4 structure (Sharov *et al*, 2017; Wang *et al*, 2018; Cheung & Díaz-Santín, 2018).

**Figure 3:**
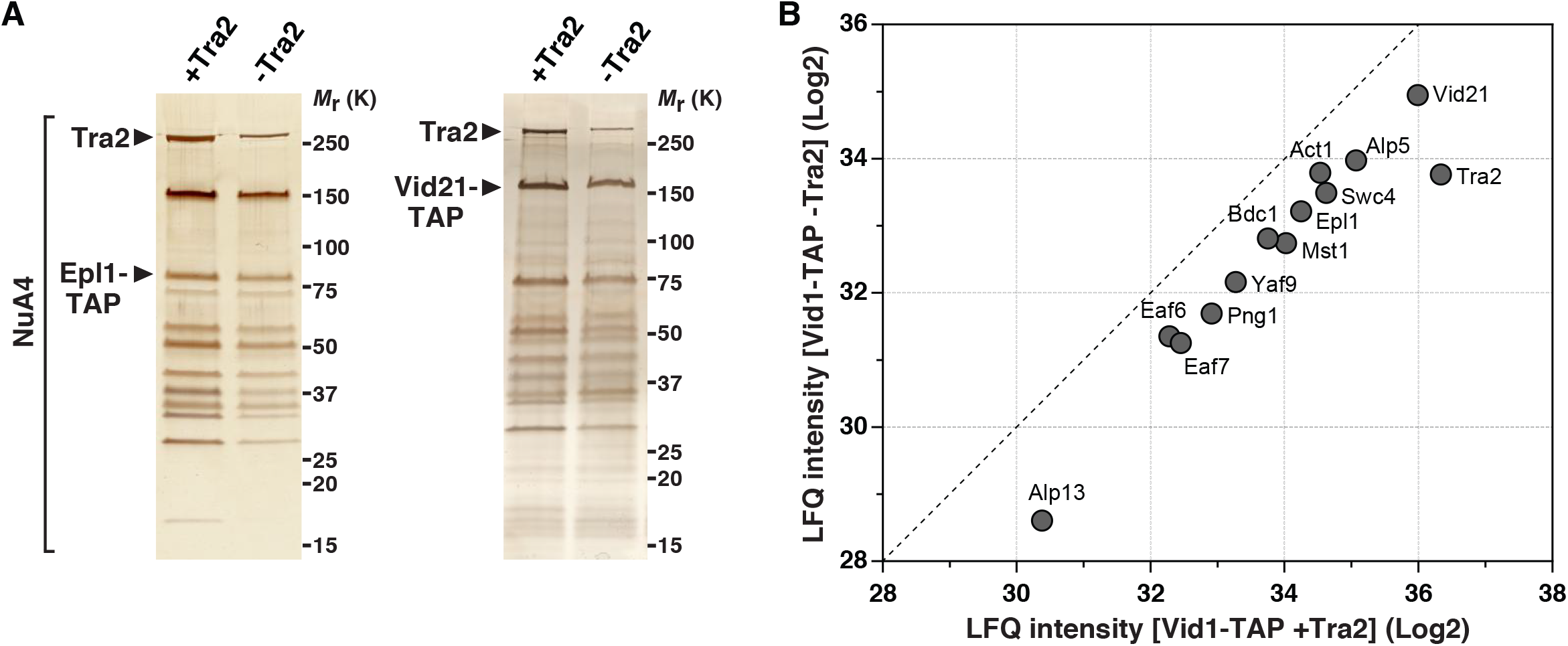
Tra2 is important for the formation of the entire NuA4 complex. (A) Silver staining of NuA4 complexes purified either in presence or absence of Tra2, using Epl1-TAP (left) or Vid21-TAP (right) as baits. *epl1-TAP tra2-CKO* and *vid21-TAP tra2-CKO* cells were grown to exponential phase in rich medium supplemented with either DMSO (+Tra2) or β-estradiol (-Tra2). (B) A scatter plot representing the LFQ intensities from LC-MS/MS analysis of NuA4 complexes purified in the presence (x-axis) or absence (y-axis) of Tra2. Individual points represent individual NuA4 subunits in Vid21-TAP eluates. The dashed line shows a 1:1 ratio.

### Mechanism for Tra1 specific interaction with SAGA

We next sought to determine how Tra1 interacts specifically with SAGA, taking advantage of the viability of *tra1* mutants in *S. pombe* and guided by the most recent cryo-electron microscopy structure of *Pichia pastoris* SAGA (Figure 4A) (Sharov *et al*, 2017). Resolution of the secondary structure elements of Tra1 bound to SAGA identified a narrow and highly flexible hinge region that was suggested to form the major, if not the single interaction surface between Tra1 and the rest of the complex. This region is located near the start of Tra1 FAT domain and consists of about 50 residues that fold into 3 distinct α-helices (H1-H3, Figure 4A). Multiple alignments of Tra1 orthologs from yeast, invertebrate, and vertebrate species indicate that this region is conserved throughout eukaryotes (Figure 4A). Interestingly, the homologous region of *S. pombe* Tra2, which is only present in NuA4, is more divergent suggesting that this region might define SAGA binding specificity.

**Figure 4:**
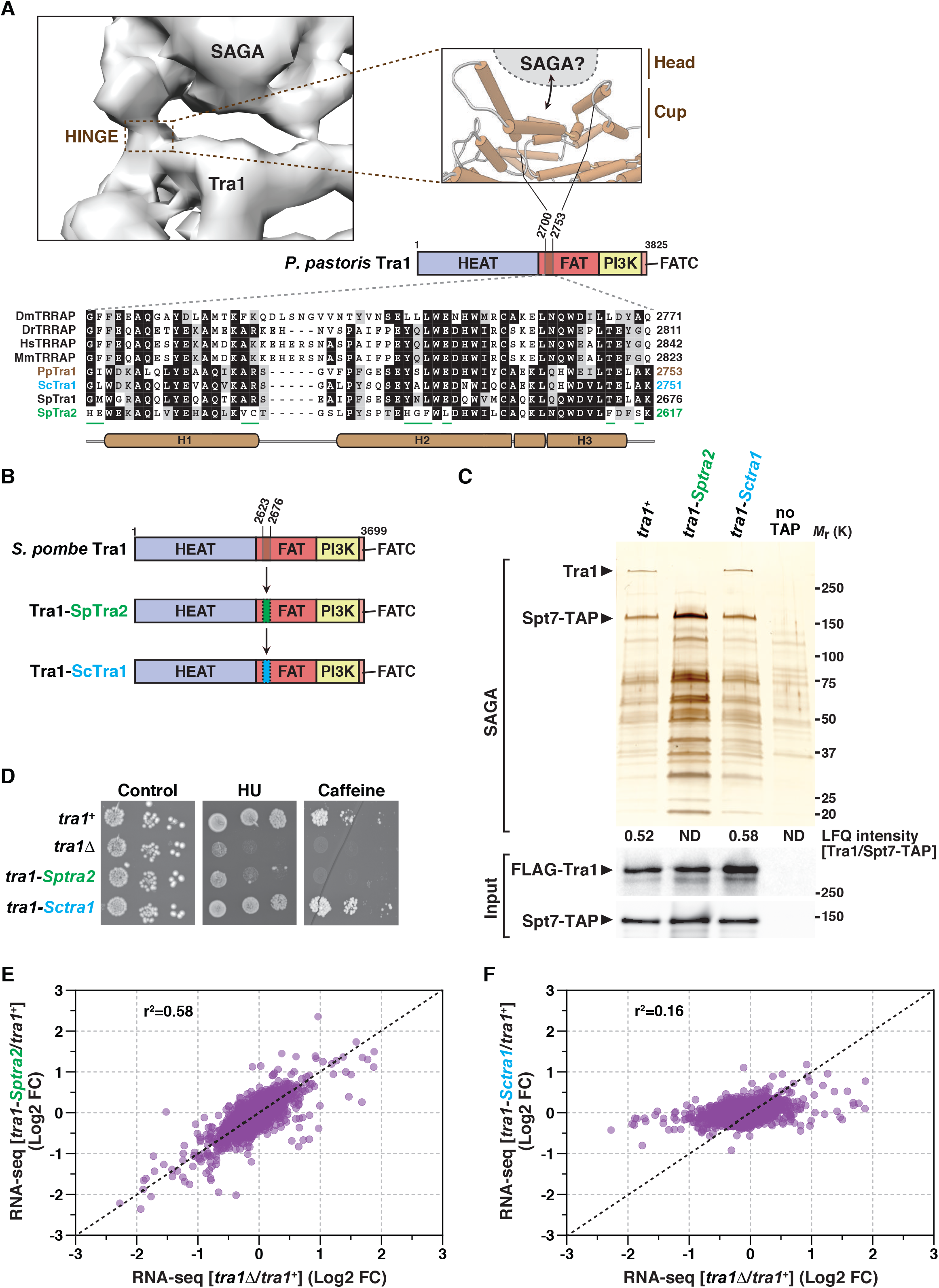
Mechanism of Tra1 specific interaction with SAGA. (A) Close-up view of the putative region of Tra1 that contacts the rest of SAGA, which constitutes the flexible hinge in the structure of *Pichia pastoris* SAGA (EMD: 3804). Cartoon cylinders represent α-helices in *P. Pastoris* Tra1 structure (PDB: 5OEJ). This domain is located near the start of the FAT domain and corresponds to residues 2700-2753 (brown-coloured box). A homologous region, defined as the <u>C</u>up <u>S</u>AGA Interacting (CSI), was identified in *S. pombe* Tra1 (residues 2623-2676) from multiple alignments of Tra1 orthologs, shown at the bottom. Residues that appear unique to *S. pombe* Tra2 are underlined (green). (B) Schematic illustration of the hybrid mutant alleles of *S. pombe tra1*+ that were constructed. Residues 2623-2676 from *S. pombe* Tra1 were swapped with the homologous region from either *S. pombe* Tra2 (green, residues 2564-2617), to create the *tra1-Sptra2* allele, or *S. cerevisiae* Tra1 (blue, residues 2698-2751), to create the *tra1-Sctra1* allele. (C) Silver staining of SAGA complexes purified from WT, *tra1-Sptra2*, and *tra1-Sctra1* strains (see B), using Spt7 as the bait. A non-tagged strain (no TAP) was used as a control for background. Numbers at the bottom of the gel represent LFQ intensity ratios of Tra1 to the bait, Spt7, from LC-MS/MS analyses of purified SAGA complexes (ND: not detected). Below are anti-FLAG and anti-HA Western blotting of FLAG-Tra1 and Spt7-TAP in a fraction of the input used for TAP. Shown are gels that are representative of five independent experiments. (D-E) HU sensitivity (D) and gene expression changes (E,F) of *tra1-Sptra2, tra1-Sctra1*, as compared to *tra1*Δ mutants. (D) Ten-fold serial dilutions of exponentially growing cells of the indicated genotypes were spotted on rich medium (control), medium supplemented with 10 mM HU, or 15 mM caffeine, and incubated at 32°C. (E,F) Scatter plots from RNA-seq count data comparing *tra1*Δ mutants (x-axis) with either *tra1-Sptra2* mutants (y-axis in E) or *tra1-Sctra1* mutants (y-axis in F), relative to isogenic WT controls. Statistical significance and correlation were analyzed by computing the Pearson correlation coefficient (r^2^ = 0.58 for *tra1-Sptra2 vs*. *tra1*Δ and r^2^ = 0.16 for *tra1-Sctra1* vs. *tra1*Δ; *P* < 0.001). The black dashed line represents a 1:1 ratio.

Deletion of a few helices within Tra1 might cause important structural rearrangements and destabilize the protein (Knutson & Hahn, 2011). Thus, to determine the contribution of this 50-residue region to Tra1-SAGA interaction, we swapped them with those from Tra1 closest homolog, *S. pombe* Tra2, which is not present in SAGA. We also introduced the corresponding sequence from *S. cerevisiae* Tra1 (Figure 4B), which is shared between SAGA and NuA4. Both Tra1-ScTra1 and Tra1-SpTra2 mutant proteins were expressed at levels similar to those of wild-type Tra1 (Figure 4C). In marked contrast, silver staining and quantitative MS analyses revealed that the Tra1-SpTra2 hybrid is not detectable in Spt7 purifications, whereas normal levels of the Tra1-ScTra1 hybrid were observed (Figure 4C). Similarly, a Tra1-mTOR hybrid protein is unable to co-purify with SAGA (Figure S7A), consistent with human mTOR assembling into different PIKK-containing complexes, TORC1 and TORC2. Importantly, both Tra1-ScTra1 and Tra1-SpTra2 hybrid proteins efficiently copurified with Tti2, as shown by quantitative MS analyses (Figure S7B). Thus, this region does not affect Tra1 binding to TTT and Tra1-SpTra2 is normally recognized by its cochaperone despite being defective in SAGA incorporation.

Phenotypic analyses of *tra1-Sctra1* and *tra1-Sptra2* strains revealed that *tra1-Sptra2* mutants are sensitive to replicative stress and caffeine, similar to *tra1*Δ mutants, whereas *tra1-Sctra1* strains showed no growth defects, as compared to wild-type cells (Figure 4D). RNA-seq analyses of *tra1-Sctra1* and *tra1-Sptra2* mutants revealed a positive correlation between the transcriptomic changes observed in *tra1-Sptra2* and *tra1*Δ mutants, as compared to a wild-type control (r^2^ = 0.58) (Figure 4E). In contrast, *tra1-Sctra1* mutants showed few gene expression changes, which correlated poorly with those observed in *tra1*Δ mutants (r^2^ = 0.16) (Figure 4F). Thus, a 50-residue region from *S. cerevisiae* Tra1 complements that of *S. pombe* Tra1, likely because *S. cerevisiae* Tra1 is present in both SAGA and NuA4. In contrast, the homologous *S. pombe* Tra2 region is more divergent and does not complement that of *S. pombe* Tra1 because Tra2 binds NuA4 specifically, possibly through other regions. Altogether, structural, biochemical and functional evidence demonstrate that Tra1 directly contacts SAGA through a restricted, 50-residue region located at the beginning the FAT domain. This region of Tra1 consists of 3 α-helices that fold into a cup-shaped structure (Figure 4A) (Sharov *et al*, 2017). We thus coined this part of the Tra1-SAGA hinge as the Cup SAGA Interacting (CSI) region of Tra1.

### The SAGA subunit Spt20 anchors Tra1 into the SAGA complex

Patrick Schultz’s laboratory reported that the hinge accommodates a putative α-helix belonging to a SAGA subunit other than Tra1 (Sharov *et al*, 2017). This observation encouraged us to identify the residues that forms the head part of the hinge and directly contacts Tra1 CSI region. Besides Tra1, there are 18 subunits in *S. pombe* SAGA (Helmlinger *et al*, 2008). Genetic, biochemical, and structural evidence suggest that, of these, Ada1, Taf12, and Spt20 are good candidates to anchor Tra1 within SAGA (Wu & Winston, 2002; Lee *et al*, 2011; Han *et al*, 2014; Setiaputra *et al*, 2015; Sharov *et al*, 2017; Helmlinger *et al*, 2011). Silver staining analyses revealed that Tra1 is undetectable in SAGA purified from *spt20*Δ mutants, without any other visible changes in its overall migration profile (Figure 5A). *S. pombe* Spt20 is thus essential for Tra1 to interact with SAGA.

**Figure 5:**
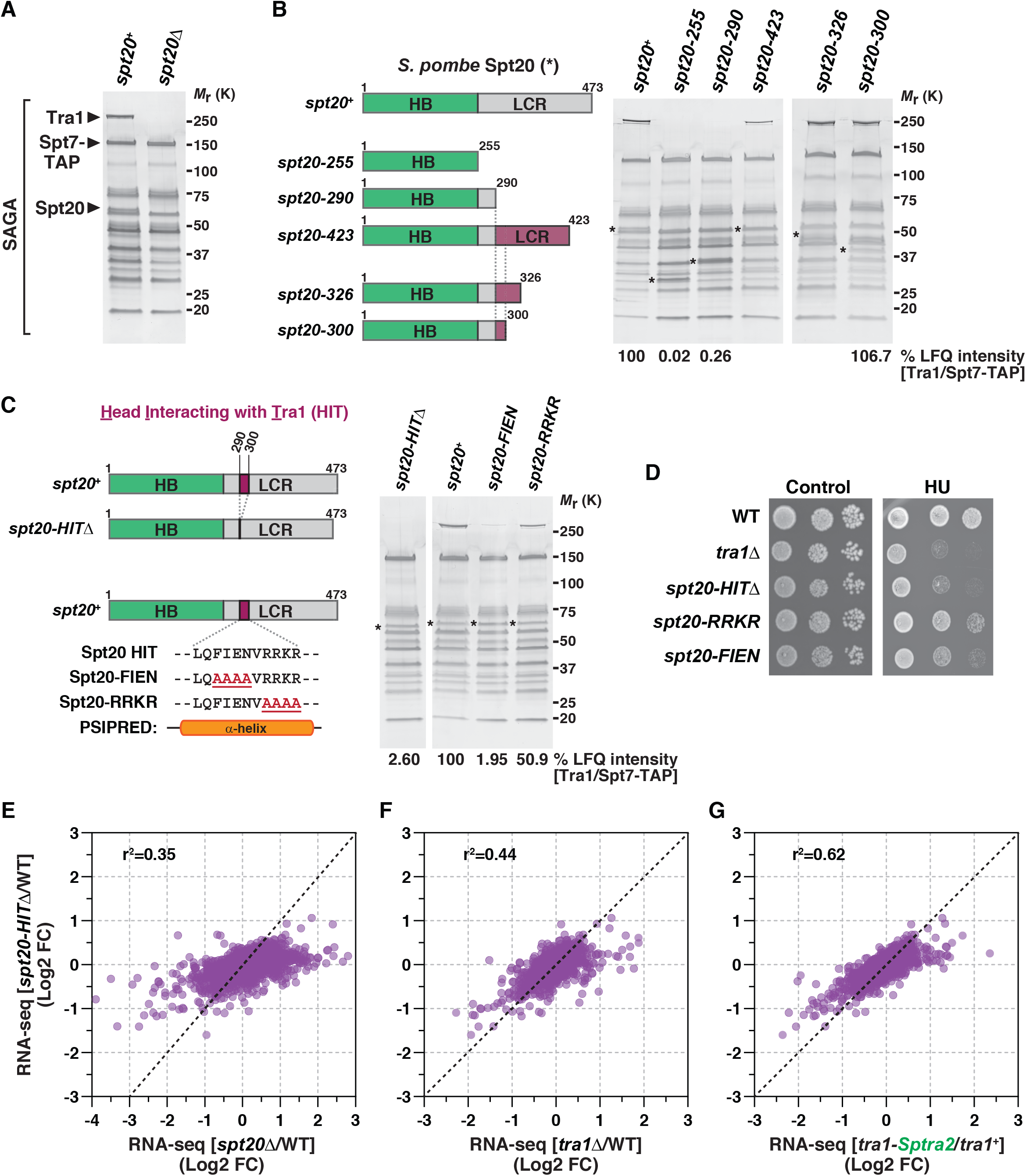
Spt20 anchors Tra1 into the SAGA complex. (A) Silver staining of SAGA complexes purified from WT and *spt20*Δ strains, using Spt7 as the bait. A band corresponding to Spt20 disappears in *spt20*Δ mutants. (B) Schematic illustration of the different Spt20 truncation mutant alleles constructed to identify the <u>H</u>ead Interacting with <u>T</u>ra1 (HIT) region of Spt20 from *S. pombe*. Distinct colours depict Spt20 domains, defined as <u>H</u>omology <u>B</u>oxes (HB) and a <u>L</u>ow <u>C</u>omplexity <u>R</u>egion (LCR). Each allele is named after to the last residue present in the truncation mutant, which shortens the LCR to various extent, as illustrated. Silver staining of SAGA complexes purified from WT and *spt20* truncation mutants, using Spt7 as the bait. Numbers at the bottom of the gel represent LFQ intensity ratios of Tra1 to Spt7, from LC-MS/MS analyses of SAGA purifications. Values for each mutant are expressed as percentage of WT SAGA. Purple colouring depicts the region of Spt20 that allows SAGA to interact with Tra1 in each truncation mutant. (C) Schematic illustration of the deletion or point mutations within Spt20 HIT region, narrowed down to residues 290-300 (purple-coloured box) and predicted to fold into a α-helix, using PSI-blast based secondary structure PREDiction (PSIPRED, (Buchan *et al*, 2013)). Silver staining of SAGA complexes purified from WT, *spt20-HIT*Δ, s*pt20-FIEN*, and *spt20-RRKR* mutants, using Spt7 as the bait. Numbers at the bottom of the gel represent LFQ intensity ratios of Tra1 to Spt7, from LC-MS/MS analyses of purified SAGA complexes. Values for each mutant are expressed as percentage of WT SAGA. (B,C) Asterisks indicate the position of the WT and mutant Spt20 proteins in purified SAGA. (D) HU sensitivity of *spt20-HIT*Δ, s*pt20-FIEN*, and *spt20-RRKR* mutants, as compared to *tra1*Δ mutant strains. Ten-fold serial dilutions of exponentially growing cells of the indicated genotypes were spotted either on rich medium (control) or medium supplemented with 5 mM HU and incubated for 3 days at 32°C. (E-G) Scatter plots from RNA-seq data comparing *spt20-HITD*Δ mutants (y-axis) with either *spt20*Δ mutants (x-axis in E), *tra1*Δ mutants (x-axis in F), or *tra1-Sptra2* mutants (x-axis in G), relative to isogenic WT controls. Statistical significance and correlation were analyzed by computing the Pearson correlation coefficient (r^2^ = 0.35 for *spt20-HIT*Δ *vs. spt20*Δ; r^2^ = 0.44 for *spt20-HIT*Δ *vs*. *tra1*Δ; r^2^ = 0.62 for *spt20-HIT*Δ *vs. tra1-Sptra2*; *P* < 0.001). The black dashed line represents a 1:1 ratio.

In *S. pombe*, Spt20 is 474 residue long and can be divided into an N-terminal half that contains several conserved regions, named homology boxes (HB) (Nagy *et al*, 2009) and a C-terminal low-complexity region (LCR) (Figure 5B). Deletion of Spt20 N-terminal half (residues 1-255) abolished its interaction with SAGA (data not shown), indicating that this portion of Spt20 mediates its binding to the complex. Silver staining analyses of SAGA purified from mutants that remove various lengths of Spt20 C-terminal LCR identified a short region of 11 residues that is crucial to incorporate Tra1 into SAGA (Figure 5B). Quantitative MS analyses confirmed that Tra1 does not interact with SAGA in *spt20-290* mutants, in which residues 291-474 are deleted, whereas normal levels of Tra1 are detected in *spt20-300* mutants, in which residues 301-474 are deleted (Figure 5B). Structure prediction of *S. pombe* Spt20 identified a α-helix in this region, which we coined the Head Interacting with Tra1 (HIT) (Figure 5C). Silver staining and quantitative MS analyses of SAGA purified from mutants in which Spt20 HIT region is deleted (*spt20-HIT*Δ) confirmed the importance of this region for Tra1 interaction (Figure 5C). Similarly, mutational analyses of the HIT identified 4 residues, FIEN, that are important for Tra1 incorporation into SAGA, whereas the next 4, positively charged RRKR residues contribute less (Figure 5C). All Spt20 truncation, deletion, and point mutants were expressed at levels comparable to those of wild-type Spt20 (Figure S8) and, importantly, were present in purified SAGA complexes (* in Figure 5B,C).

We next evaluated the phenotype of *spt20-HIT* mutant strains. Similar to *tra1*Δ mutants, *spt20-HIT*Δ and *spt20-FIEN* mutants showed sensitivity to replicative stress, whereas *spt20-RRKR* showed milder defects, as compared to wild-type cells (Figure 5D). We then performed RNA-seq analyses of *spt20*Δ and *spt20-HIT*Δ mutants and paralleled the transcriptomic changes with those observed in *tra1-Sptra2* and *tra1*Δ mutants, as compared to a wild-type control strain. First, comparing *spt20-HIT*Δ with *spt20*Δ mutants revealed that Spt20 HIT region contributes to the expression of only a subset of Spt20-dependent genes (r^2^ = 0.35) (Figure 5E), consistent with the HIT region being specifically involved in Tra1 interaction. Indeed, comparing *spt20-HIT*Δ with *tra1*Δ mutants resulted in a better correlation (r^2^ = 0.44) (Figure 5F). Remarkably, the best correlation was obtained when comparing *spt20-HIT*Δ with *tra1-Sptra2* mutants (r^2^ = 0.62) (Figure 5G), *ie* strains in which the hinge is mutated on either side of the same interaction surface. Altogether, biochemical and functional approaches identified a narrow region of Spt20 which is necessary to incorporate Tra1 into SAGA, likely by direct interaction with Tra1 CSI region.

### Spt20 is sufficient to incorporate Tra1 specifically into SAGA in both *S. pombe* and *S. cerevisiae*

We next considered that such a restricted and specific interaction surface has appeared in *S. pombe* because Tra1 and Tra2 have diverged enough to interact exclusively with either SAGA or NuA4, respectively. To test this possibility, we sought to identify Spt20 HIT region in the budding yeast *S. cerevisiae*, which single Tra1 protein interacts with both SAGA and NuA4.

Although *S. pombe* and *S. cerevisiae* Spt20 orthologs diverge substantially, their overall domain organization remain similar. We therefore focused our mutational analysis of *S. cerevisiae* Spt20 interaction with Tra1 to a region located between the HB and LCR domains (Figure 6A). Silver staining analyses of *S. cerevisiae* SAGA purified from mutants that remove various lengths of Spt20 C-terminal LCR identified a short region of 18 residues that is critical to incorporate Tra1 into SAGA (Figure 6A). Western blotting analyses of Spt20 truncation mutants confirmed that each mutant is still present in SAGA, suggesting that this region of Spt20 is directly involved in Tra1 interaction (Figure 6B). Structure prediction of *S. cerevisiae* Spt20 identified a α-helix in this region (Figure 6A), suggesting that Spt20 HIT region is functionally and structurally conserved between *S. pombe* and *S. cerevisiae*.

**Figure 6:**
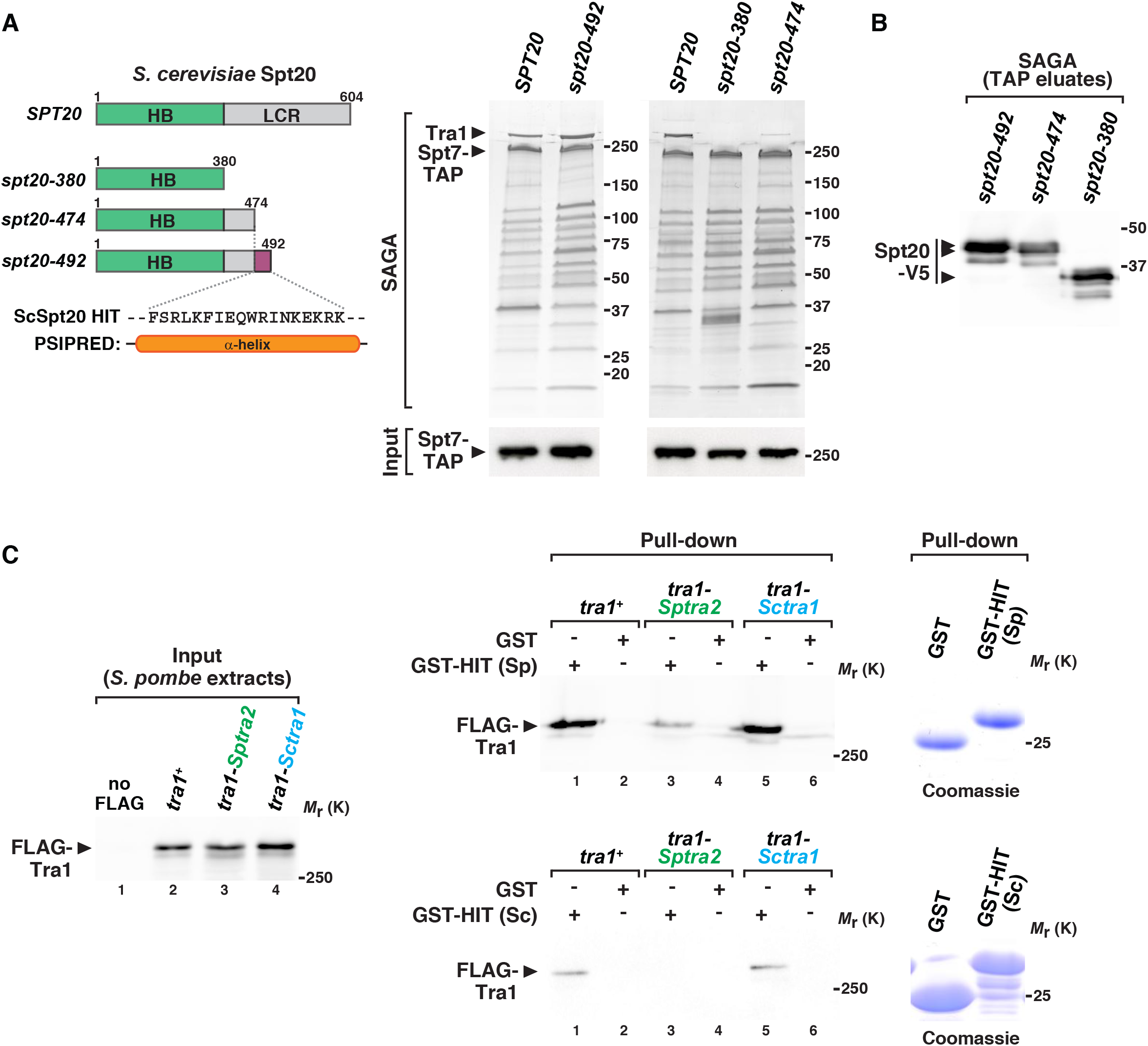
Spt20 HIT region is conserved and sufficient to incorporate Tra1 specifically into SAGA. (A) Schematic illustration of the different Spt20 truncation mutant alleles constructed to identify the <u>H</u>ead Interacting with <u>T</u>ra1 (HIT) region of Spt20 from *S. cerevisiae*. Distinct colours depict Spt20 domains, defined as <u>H</u>omology <u>B</u>oxes (HB) and a <u>L</u>ow <u>C</u>omplexity <u>R</u>egion (LCR). Each allele is named after to the last residue present in the truncation mutant, which shortens the LCR to various extent, as illustrated. Silver staining of *S. cerevisiae* SAGA complexes purified from WT and *spt20* truncation mutants, using Spt7 as the bait. Below is an anti-HA Western blot of HA-Spt7-TAP in a fraction of the input used for TAP. Shown are gels that are representative of two independent experiments. In *S. cerevisiae*, the HIT region of Spt20 is narrowed down to residues 474-492 (purple-coloured box) and predicted to fold into a α-helix, using PSI-blast based secondary structure PREDiction (PSIPRED, (Buchan *et al*, 2013)). (B) Western blot analyses of V5-tagged Spt20 truncation mutants in SAGA complex eluates, tandem affinity purified as in (A). (C) GST pull-down of *S. pombe* protein extracts from WT, *tra1-Sptra2*, and *tra1-Sctra1* strains, using GST fused to *S. pombe* (Sp) Spt20 HIT region (residues 282-324) (top right panel) or GST fused to *S. cerevisiae* (Sc) Spt20 HIT region (residues 468-537) (bottom right panel). GST alone serves as a negative control for background binding to glutathione beads. Anti-FLAG Western blotting is used to detect WT and hybrid Tra1 proteins bound to GST or GST-HIT columns (right panel) and in a fraction (0.6%) of the input used for the pull-downs. Shown are gels that are representative of three independent experiments. Coomassie blue staining of purified GST fusion proteins are shown on the right.

Finally, we asked whether Spt20 HIT region is sufficient to interact with Tra1. For this, a peptide of about 50 residues encompassing the HIT region from either *S. pombe* or *S. cerevisiae* was immobilized, through fusion to GST, and incubated with *S. pombe* protein extracts prepared from wild-type, *tra1-Sptra2* and *tra1-Sctra1* strains. Both the *S. pombe* and *S. cerevisiae* Spt20 HIT protein fragments specifically pulled down wild-type *S. pombe* Tra1 and the Tra1-ScTra1 hybrid proteins, as compared to GST alone (lanes 1 *vs* 2 and 5 *vs* 6, Figure 6C). Consistent with *in vivo* observations that Tra1 CSI region mediates interaction with Spt20, lower amounts of the Tra1-SpTra2 hybrid protein were recovered on the GST-HIT column, (lane 3 *vs* 4, Figure 6C). Overall, these experiments indicate that Spt20 HIT region folds into a α-helix that is both necessary and sufficient for binding to Tra1 in both *S. pombe* and *S. cerevisiae*. We have thus deciphered the molecular topology of the narrow hinge that mediates the specific contact between Tra1 and the rest of SAGA. Remarkably, only a few residues are involved on each side of the SAGA-Tra1 interface, in agreement with the peripheral position of Tra1 within SAGA (Sharov *et al*, 2017).

### Tra1 orchestrates an ordered pathway for SAGA assembly

Throughout this study, quantitative MS analyses of *S. pombe* SAGA purified from various mutants revealed an unexpected finding. Indeed, the amount of DUB module subunits within SAGA consistently decreased when Tra1 was not incorporated into the complex. For instance, we measured a reproducible decrease of both Sgf73 and Ubp8 in Spt7 purified from *tra1*Δ or β-estradiol-treated *tti2-CKO* cells, as compared to control conditions (Figure 7A,B). Similarly, mutating either side of the hinge reduced the amount of both Sgf73 and Ubp8 in SAGA purifications, as shown in *spt20-290, spt20-HIT*Δ, *spt20-FIEN*, and *tra1-Sptra2* cells (‘Hinge*’ in Figure 7A,B). In contrast, the levels of Sgf73 and Ubp8 did not change in Spt7 purifications from *spt20-300, spt20-RRKR* and *tra1-Sctra1* cells, in which Tra1 incorporates into SAGA (data not shown). The other two DUB subunits, Sgf11 and Sus1, are about 10 kDa and therefore less reliably quantified by MS. Still, the reproducibility of this effect across distinct mutants that all affect Tra1 incorporation suggested that Tra1 promotes the assembly of the DUB module into SAGA.

**Figure 7:**
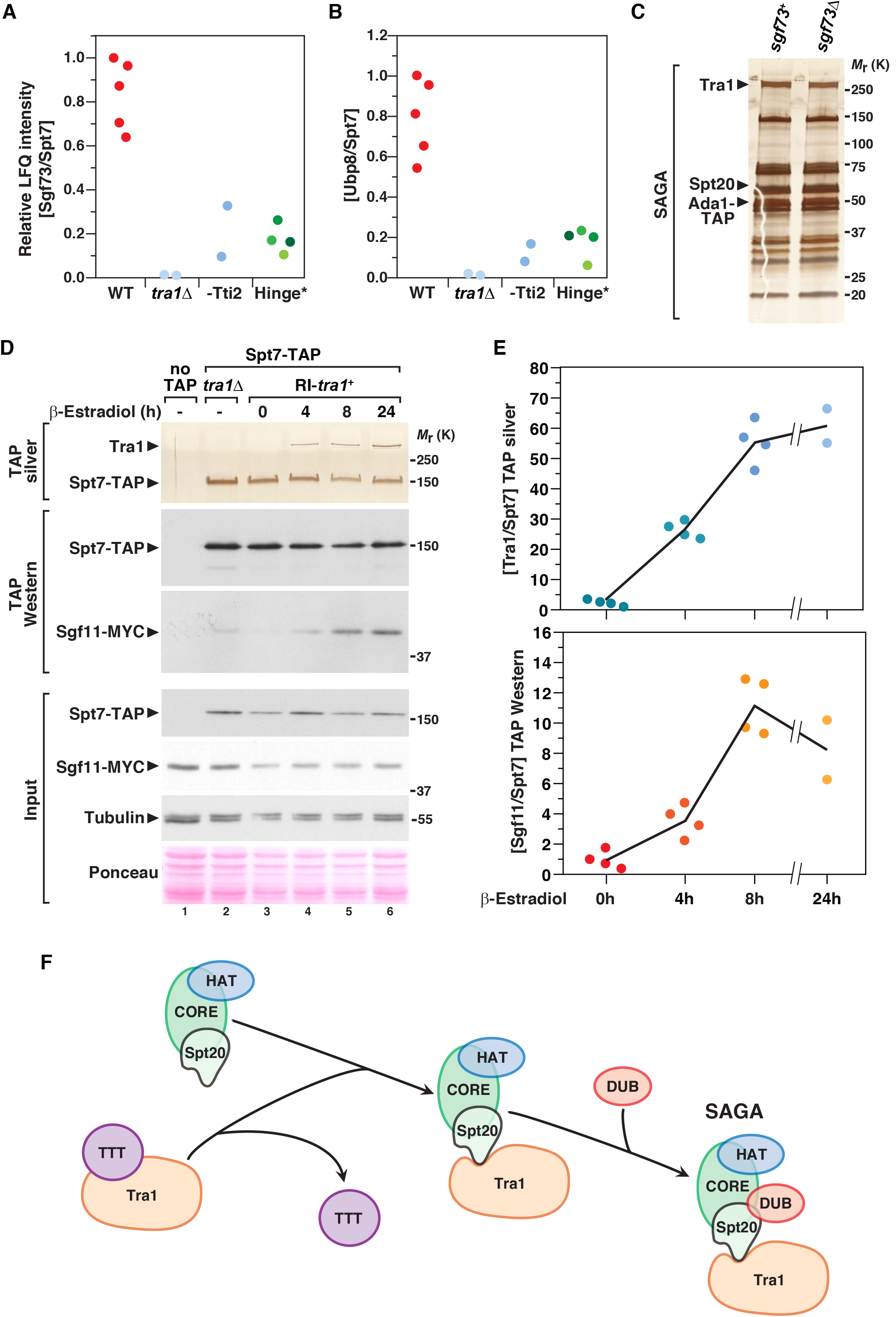
Tra1 promotes the incorporation of the DUB module within SAGA. (A,B) LC-MS/MS analysis of SAGA complexes purified from mutants defective in Tra1-SAGA interaction, including *tra1*Δ (n = 2) and β-estradiol treated *tti2-CKO* mutants (-Tti2, n = 2), as well as four distinct mutant alleles that disrupt the hinge region (Hinge*). These include *tra1-Sptra2, spt20-290, spt20-HIT*Δ, and *spt20-FIEN* mutants, labelled from dark to light green, respectively. Relative LFQ intensity ratios of Sgf73 (A) and Ubp8 (B) to the bait, Spt7, from independent experiments or mutants are plotted individually. (C) Silver staining of SAGA complexes purified from WT (*sgf73*+) or *sgf73*Δ mutants, using Ada1 as the bait. Shown are gels that are representative of three independent experiments. (D) Silver staining and Western blotting of SAGA complexes purified upon Tra1 *de novo* expression from a strain in which the DUB subunit Sgf11 is MYC-tagged. *spt7-TAP RI-tra1 sgf11-MYC* cells were grown to exponential phase and harvested at different time-points after β-estradiol addition, as indicated (hours). SAGA was purified from a *tra1*Δ strain as a control for the complete loss of Tra1 from SAGA and from a non-tagged strain (no TAP) as a control for background. Silver staining reveals Spt7 and Tra1, which migrate around 150 and 400 kDa, respectively. Anti-HA and anti-MYC Western blotting of Spt7-TAP and Sgf11-MYC in a fraction of the input (Input) and in TAP eluates (TAP) is shown below. An anti-tubulin antibody and Ponceau red staining are used as loading controls. Shown are gels that are representative of four independent experiments, quantified and averaged in (E). (E) Quantification of the ratio of Tra1 to Spt7 from silver stained gels (top) and of the ratio of Sgf11-MYC to Spt7-TAP from Western blots (bottom). Data points were individually plotted on the graph. Signal intensities were quantified from 2-4 independent experiments. (F) Working model for the last steps of SAGA assembly. Core subunits (Spt7, Ada1, and TAFs), the HAT module (Gcn5, Ada2, Ada3, and Sgf29), and Spt20 form a pre-assembled complex. The Hsp90 cochaperone TTT promotes the maturation of nascent Tra1, which is then anchored to SAGA by the HIT domain from Spt20. Consequently, Tra1 stabilizes the interaction of the DUB (Sgf73, Ubp8, Sgf11, and Sus1) with SAGA to form a mature, fully active multifunctional transcriptional co-activator complex.

We first asked whether, conversely, the DUB stabilizes Tra1 within SAGA. In *S. cerevisiae*, Sgf73 is critical to anchor the DUB module into SAGA (Köhler *et al*, 2008). Mass spectrometry analyses confirmed that, in *S. pombe sgf73*Δ mutants, the DUB subunits Ubp8, Sgf11, and Sus1 were absent from SAGA purifications (data not shown). In contrast, silver staining and MS analyses indicated that Spt20 and Tra1 incorporation into SAGA was similar between wild-type and *sgf73*Δ strains (Figure 7C). Altogether, these results suggest that SAGA assembly follows a directional, ordered pathway, in which Spt20 anchors Tra1, which then stabilizes the DUB into SAGA.

To test this hypothesis directly, we purified SAGA from *RI-tra1+* strains (Figure 2C and Figure S4), in which we tagged endogenous Sgf11 using a MYC epitope. We then concomitantly monitored the kinetics of the *de novo* incorporation of Tra1 and the DUB module into SAGA. First, we confirmed that Sgf11 interacts less strongly with Spt7 in *tra1*Δ mutants or untreated *RI-tra1*+ cells, as compared to control conditions (lanes 2 and 3 *vs* 6, Figure 7D). Second, we detected a progressive increase in the amount of Sgf11 that interacts with Spt7 upon β-estradiol treatment and *de novo* incorporation of Tra1 into SAGA (lanes 4 and 5, Figure 7D). Quantification of independent experiments confirmed these observations (Figure 7E). Comparing the relative levels of Sgf11 and Tra1 in SAGA at 4 hours suggested that assembly of the DUB module is slightly delayed, as compared to Tra1.

Overall, we accumulated functional and biochemical evidence that support a model in which nascent Tra1 is bound by the TTT cochaperone, possibly to promote its folding into a mature conformation. Tra1 is then assembled by direct interaction with a narrow region of Spt20 and then promotes the incorporation of the DUB module within SAGA (Figure 7F).

## Discussion

Many chromatin and transcription regulators function within large multimeric complexes. Deciphering the principles that govern their assembly is key to understand their structural organization, function, and regulation. Our work brings several mechanistic insights into the *de novo* assembly and modular organization of two such complexes, the SAGA and NuA4 transcription co-activators. First, functional and biochemical evidence indicate that the Hsp90 cochaperone TTT promotes Tra1 and Tra2 incorporation into SAGA and NuA4, respectively. Second, structure-guided mutational analyses elucidated the specificity of Tra1 interaction with SAGA *versus* NuA4. We further defined the topology of the Tra1-SAGA interaction surface, which, remarkably, appears restricted to a small region of Tra1 FAT domain contacting a single α-helix of the core subunit Spt20. Third, in contrast to the general role of Tra2 in NuA4 complex formation, Tra1 specifically controls the incorporation of the DUB module into SAGA, uncovering an ordered pathway of SAGA assembly (Figure 7F).

### Chaperone-mediated assembly of the SAGA and NuA4 co-activator complexes

Our work indicates that SAGA and NuA4 require, at least in part, a dedicated chaperone machinery for their assembly. A recent study showed that human TAF5 and its paralog TAF5L, which are specific to the general transcription factor TFIID and SAGA, respectively, require a dedicated chaperone, the CCT chaperonin, for incorporation into pre-assembled modules (Antonova *et al*, 2018). Alternatively, cotranslational assembly has appeared as a prevalent regulatory mechanism for promoting protein-protein interactions in eukaryotes (Duncan & Mata, 2011; Shiber *et al*, 2018). Cotranslational interactions were observed for subunits of the SET1C histone methyltransferase complex (Halbach et al, 2009) and, more recently, between specific SAGA subunits in *S. cerevisiae* and human cells (Kassem et al, 2017; Kamenova et al, 2018). Our study therefore contributes to the emerging concept that dedicated chaperone machineries and ordered pathways control the *de novo* assembly of chromatin and transcription regulatory complexes.

### Evolutionary conservation of the Tra1 pseudokinase in the PIKK family

Studies in mammals revealed that the pleiotropic HSP90 chaperone is specifically recruited to PIKKs by a dedicated cochaperone, the TTT complex, to promote their stabilization and incorporation into active complexes (Takai *et al*, 2007; Anderson *et al*, 2008; Takai *et al*, 2010; Hurov *et al*, 2010; Kaizuka *et al*, 2010; Izumi *et al*, 2012). In contrast, the effect of TTT on the Tra1 pseudokinase, the only inactive PIKK, is less characterized. Previous work showed that TTT stabilizes TRRAP in human cells (Takai *et al*, 2007; Kaizuka *et al*, 2010; Hurov *et al*, 2010; Izumi *et al*, 2012) and several studies reported physical and genetic interaction between Tra1 and TTT components in yeast (Hayashi *et al*, 2007; Shevchenko *et al*, 2008; Helmlinger *et al*, 2011; Genereaux *et al*, 2012; Inoue *et al*, 2017).

We accumulated functional and biochemical evidence that, in *S. pombe*, Hsp90 and TTT promote the incorporation of Tra1 and Tra2 into SAGA and NuA4 complexes, respectively. In agreement, we found that the TTT subunit Tti2 contributes to Tra1- and Tra2-dependent gene expression, as well as SAGA and NuA4 promoter recruitment. Therefore, although Tra1 is the sole catalytically inactive member of the PIKK family, it shares a dedicated chaperone machinery with active PIKK kinases for its folding, maturation, and assembly into a larger complex.

Phylogenetic analyses of PIKK orthologs in various organisms indicate that the Tra1 pseudokinase appeared early in the eukaryotic lineage, concomitantly with other PIKKs (our unpublished observations). As expected for a pseudokinase, the catalytic residues diverge substantially. However, Tra1 orthologs show high conservation of PIKK distinctive domain architecture, which consists of a long stretch of helical HEAT repeats, followed by TPR repeats forming the FAT domain, preceding FRB, PI3K-like, and FATC domains. It is thus tempting to speculate that the requirement of PIKKs for a dedicated cochaperone explains the selection pressure that is observed on the sequence and domain organization of Tra1, in the absence of conserved, functional catalytic residues. For example, the short, highly conserved C-terminal FATC domain loops back close to the active site and is critical for mTOR kinase activity (Imseng *et al*, 2018). Similarly, we found that the FATC is essential for Tra1 incorporation into SAGA (our unpublished observations), presumably through allosteric control of the folding and positioning of Tra1 CSI region, which directly contacts SAGA.

### Distinct architectural roles of Tra1 between the SAGA and NuA4 complexes

Biochemical and functional evidence suggested that the Tra1 pseudokinase serves as a scaffold for the assembly and recruitment of the SAGA and NuA4 complexes to chromatin. *S. pombe* provides a unique opportunity to better understand its roles within each complex because it has two paralogous proteins, Tra1 and Tra2, and each has non-redundant roles that are specific for SAGA or NuA4, respectively (Helmlinger *et al*, 2011).

Our work establishes that, within SAGA, Tra1 has specific regulatory roles and does not scaffold the entire complex but, rather, controls the assembly of the DUB module. In contrast, Tra2 appears to contribute to the structural integrity of NuA4. In agreement, biochemical and structural analyses of yeast SAGA and NuA4 complexes reveal distinct positioning of Tra1 relative to other components. Within SAGA, Tra1 localizes to the periphery of the SAGA complex (Sharov *et al*, 2017) and directly interacts with Spt20 (Figure 5), whereas it occupies a more central position within NuA4 and contacts several different subunits (Wang *et al*, 2018). We therefore anticipate that the single Tra1 protein found in most other eukaryotic organisms will have distinct structural roles between SAGA and NuA4 and function as a scaffold only for the NuA4 complex.

### Topological organization of the Tra1-SAGA interface

In marked contrast with *S. cerevisiae* and mammals, a *tra1*Δ deletion mutant is viable in *S. pombe*, enabling detailed biochemical and genetic studies that are not possible in other organisms (Helmlinger, 2012). We indeed made significant progress in the characterization of Tra1 incorporation into SAGA. The latest structure of SAGA clearly shows that Tra1 occupies a peripheral position and interacts with the rest of the complex through a narrow and flexible surface interaction, forming a hinge (Sharov *et al*, 2017). Our structure-function analyses identified the residues that constitute the hinge.

Specifically, we show that a restricted, 50-residue region of the large Tra1 protein dictates the specificity of its interaction with SAGA. The homologous region from *S. pombe* Tra2 diverged such that it cannot interact with SAGA. Conversely, within the hinge, an additional density predicted to form a α-helix and not attributable to Tra1 was observed at the threshold used to resolve Tra1 secondary structure elements (Sharov *et al*, 2017). We demonstrate that a small portion of Spt20, the HIT region, is both necessary and sufficient to anchor Tra1 within SAGA. This region of Spt20 constitutes the major interaction interface between Tra1 and the rest of SAGA in both *S. pombe* and *S. cerevisiae*, allowing the construction of unique separation-of-function alleles for phenotypic and transcriptomic analyses. The exact roles of Tra1/TRRAP have been indeed challenging to study genetically because of its presence in both SAGA and NuA4 (Knutson & Hahn, 2011) and because it has essential roles in *S. cerevisiae* proliferation or during mouse early embryonic development (Saleh *et al*, 1998; Herceg *et al*, 2001). As we show using *S. cerevisiae* (Figure 6), the identification of the specific residues that mediates most, if not all, Tra1-SAGA contacts enables the design of unique mutant alleles for studying the exact roles of Tra1/TRRAP within SAGA *versus* NuA4.

Finally, these findings open new perspectives to better understand the molecular mechanism by which Tra1 modulates SAGA enzymatic activities upon binding transcription activators. Along this line, the observed structural flexibility of the hinge region might be functionally important and suggests that the interaction between Tra1 and Spt20 HIT region is highly dynamic. Understanding the molecular basis and functional relevance of this flexibility will undoubtedly be an important goal for future research projects but will likely require novel methodological approaches.

### Ordered assembly pathway of the SAGA complex

Seminal work revealed that complexes are generally assembled by ordered pathways that appear evolutionarily conserved (Marsh & Teichmann, 2015). Biochemical analyses of SAGA in various mutants suggested that the last steps of SAGA assembly occur through an ordered pathway. Indeed, Spt20 is required for both Tra1 and DUB incorporation into SAGA, while Tra1 stabilizes the DUB, but not Spt20, within SAGA. Conversely, the DUB module does not modulate Spt20 or Tra1 assembly. Finally, monitoring the fate of the DUB component Sgf11 upon Tra1 *de novo* synthesis supports a model in which Tra1 interacts with Spt20 and then stabilizes the DUB module within the complex (Figure 7F).

However, Tra1 is presumably not directly recruiting the DUB module into SAGA. Recent structural analyses indicate that Tra1 does not stably contact any DUB component in the majority of mature SAGA complexes (Sharov *et al*, 2017). Rather, Tra1 might stabilize DUB incorporation during the assembly process, either through transient, direct interaction or indirectly, by inducing a conformational change within Spt20 that promotes SAGA-DUB interactions. Combining our work with previous structural and biochemical analyses suggest that Spt20 might directly contact the DUB anchor subunit, Sgf73, although a higher resolution structure of SAGA is eventually needed to validate this hypothesis.

Overall, our findings contribute to our understanding of how multifunctional chromatin regulatory complexes are assembled, which is essential to better characterize their structural organization and functions. Tra1 mediates the trans-activation signal from promoter-bound transcription factors to SAGA and NuA4 regulatory activities, which have critical roles in both basal and inducible RNA polymerase II transcription. Our work opens exciting prospects for the characterization of SAGA and NuA4 functions during transcription.

## Materials and methods

### Yeast procedures and growth conditions

Standard culture media and genetic manipulations were used. *S. cerevisiae* strains were grown in YPD at 30°C to mid-log phase (∼1 x 10^7^ cells/ml). *S. pombe* strains were grown in either rich (YES) or minimal (EMM) media at 32°C to mid-log phase (∼0.5 x 10^7^ cells/ml). Proliferation assays were performed by inoculating single colonies in liquid media and counting the number of cells at different time points. For longer time course, cultures were diluted to keep cells in constant exponential growth. For auxin-inducible targeted protein degradation (AID), cells were grown at 25°C and treated with either 0.5 mM indol-3-acetic acid (IAA, I2886, Sigma) or ethanol. For CreER-loxP-mediated recombination, cells were treated with either 1 μM β-estradiol (E2758, Sigma) or DMSO alone.

### Strain construction

All *S. pombe* and *S. cerevisiae* strains used are listed in Table S2 and were constructed by standard procedures, using either yeast transformation or genetic crosses. Strains with gene deletions, truncations, or C-terminally epitope-tagged proteins were constructed by PCR-based gene targeting of the respective open reading frame (ORF) with *kanMX6, natMX6 or hphMX6* cassettes, amplified from pFA6a backbone plasmids (Bahler *et al*, 1998; Hentges *et al*, 2005). For insertion of loxP sites, the same resistance cassettes were amplified from the pUG6 or pUG75 plasmids (Euroscarf #P30114, and #P30671, respectively) (Gueldener *et al*, 2002). Constructions of point mutations, internal deletions, or domain swaps in *spt20*+ and *tra1*+ were performed using a *ura4* cassette in a two-step *in vivo* site-directed mutagenesis procedure (Storici *et al*, 2001). Alternatively, CRISPR-Cas9-mediated genome editing was used, as described in (Zhang *et al*, 2018), for example for marker-less N-terminal epitope tagging of *tra1*+. DNA fragments used for homologous recombination were generated by PCR, Gibson assembly cloning (kit E2611L, New England Biolabs), or gene synthesis. Cloning strategies and primers were designed using the online fission yeast database, PomBase (Lock *et al*, 2018). All primer sequences are listed in Table S3. Transformants were screened for correct integration by PCR and, when appropriate, verified by Sanger sequencing or Western blotting. For each transformation, 2-4 individual clones were purified and analyzed.

Because the *tti2*+ gene is essential for viability in *S. pombe* (Inoue *et al*, 2017), C-terminal epitope tagging was performed in diploids, to generate heterozygous alleles. Their sporulation demonstrated that all C-terminally tagged Tti2 strains grew similarly to wild-type controls in all conditions that were tested (data not shown).

### Plasmid construction

Auxin-inducible degron (AID) tagging was performed using a plasmid, DHB137, which we constructed by inserting three HA epitopes in fusion with the three copies of the mini-AID sequence from pMK151 (Kubota *et al*, 2013). V5-PK tagging was performed using a plasmid, DHB123, which we constructed by inserting three V5 epitopes 5’ to the hphMX6 cassette into pFA6a-hphMX6 (Euroscarf #P30438) (Hentges *et al*, 2005). For GST pull-down assays, DNA fragments comprising either nucleotides +925 to +1054 from the *S. pombe spt20* coding sequence (CDS), encoding residues Asp282 to Ala324, or nucleotides +1402 to +1611 from the *S. cerevisiae SPT20* CDS, encoding residues Met468 to Ala537, were synthesized and amplified. Each product was then subcloned into pGEX-4T2 (GE Healthcare Life Sciences), 3’ and in frame to the GST coding sequence, using the Gibson assembly kit (E2611L, New England Biolabs), to generate the DHB179 and DHB193 plasmids, respectively.

### RT-qPCR analysis

Reverse transcription and quantitative PCR analyses of cDNA were performed using RNA extracted from 50 mL of exponentially growing cells, as described in (Laboucarié *et al*, 2017), and according to the MIQE guidelines (Bustin *et al*, 2009). Briefly, total RNA was purified using hot, acidic phenol and contaminating DNA was removed by DNase I digestion, using the TURBO DNA-free™ kit (AM1907, Ambion). 1 μg of RNA was then reverse-transcribed (RT) at 55°C with random hexanucleotide primers, using the SuperScript III First-Strand System (18080051, ThermoFisher Scientific). Fluorescence-based quantitative PCR was performed with SYBR Green and used to calculate relative cDNA quantities, from the slope produced by standard curves for each primer pair, in each experiment. DNase-treated RNA samples were used as controls for the presence of genomic DNA contaminants. Standard curve slopes were comprised between −3.5 (90% efficiency) and −3.15 (110% efficiency), with an *r*^2^ > 0.9. All primer sequences are listed in Table S3.

### Protein extraction

Protein extracts were prepared as described in (Laboucarié *et al*, 2017). Briefly, 10 to 25 mL cultures of exponentially growing cells were homogenized by glass bead-beating in a FastPrep (MP Biomedicals). Proteins extracted using either standard lysis buffer (WEB: 40 mM HEPES-NaOH pH 7.4, 350 mM NaCl, 0.1% NP40, and 10% glycerol) or trichloroacetic acid (TCA) precipitation. WEB was supplemented with protease inhibitors, including cOmplete EDTA-free cocktails tablets (04693132001, Roche), 1 mM PMSF (P7626, Sigma), 1 μg/ml bestatin (B8385, Sigma), and 1 μg/ml pepstatin A (P5318, Sigma). Protein concentrations were measured by the Bradford method. Ponceau red or Coomassie blue staining were used to normalize for total protein levels across samples.

### Western blotting and antibodies

Western blotting was performed using the following antibodies: peroxidase-anti-peroxidase (PAP) (P1291, Sigma), anti-Calmodulin binding protein (CBP) (RCBP-45A-Z, ICLab), anti-tubulin (B-5-1-2, Sigma), anti-FLAG (M2, F1804, Sigma), anti-MYC (9E10, Agro-Bio LC; 9E11, ab56, Abcam; and rabbit polyclonal ab9106, Abcam), anti-V5 (SV5-Pk1, AbD Serotec), anti-HA (16B12, Ozyme; rabbit polyclonal, ab9110, Abcam). Protein concentrations were measured by the Bradford method and used to load equal amounts of proteins across samples. Quantification of signal intensity was performed using staining, film exposure, or digital acquisition that were within the linear range of detection, as verified by loading serial dilutions of one sample, and analyzed with Image Studio™ Lite 4.0 (LI-COR Biosciences).

### Chromatin immunoprecipitation

ChIP experiments were performed as previously described (Helmlinger *et al*, 2011). Briefly, cell cultures were crosslinked in 1% formaldehyde for 30 min. Cells were then broken using a FastPrep (MP Biomedicals), and the chromatin fraction was sheared to 200– 500 bp fragments using a Branson sonicator for 9 cycles (10 seconds ON, 50 seconds OFF) at an amplitude of 20%. For immunoprecipitation (IP), 3-5 μg of anti-HA (16B12) or anti-Myc antibodies (9E11) were incubated overnight at 4°C with the chromatin extracts and then coupled with 50 μl of protein-G-sepharose beads (GE17-0618-01, Sigma) during 4h at 4°C. ChIP DNA was quantified by fluorescence-based quantitative PCR using SYBR Green, as described for RT-qPCR analysis. Input (IN) samples were diluted 200-fold while IP samples were diluted 3-fold. Relative occupancy levels were determined by dividing the IP by the IN value (IP/IN) for each amplicon. To determine the specificity of enrichment of the tagged protein, the corresponding untagged control samples were included in each ChIP experiment. All primer sequences are listed in Table S3.

### Affinity purification

Protein complexes were purified by the tandem affinity purification (TAP) method, as described previously (Rigaut *et al*, 1999; Helmlinger *et al*, 2008), with minor modifications. 1-4 liters of exponentially growing cells were harvested, snap-frozen as individual droplets, and grinded in liquid nitrogen using a Freezer/Mill® (Spex SamplePrep). Protein extraction was performed in either WEB buffer or CHAPS-containing lysis buffer (CLB) buffer (50mM HEPES-NaOH pH 7.4, 300mM NaCl, 5mM CHAPS, 0.5mM DTT), supplemented with protease and phosphatase inhibitors. Following purifications, 10% of 2 mM EGTA eluates were concentrated and separated on 4%– 20% gradient SDS-polyacrylamide Tris-glycine gels (Biorad). Total protein content was visualized by silver staining, using the SilverQuest kit (LC6070, ThermoFisher Scientific). For quantitative mass spectrometry analyses, 90% of 2 mM EGTA eluates were precipitated with TCA and analyzed by mass spectrometry (MS). A downscaled version of the TAP procedure was used for standard co-immunoprecipitation followed by Western blot analysis, as described in (Laboucarié *et al*, 2017).

Recombinant GST and GST-HIT proteins were produced by IPTG induction of transformed BL21 Rosetta strains and purified on 100 μl of Glutathione Sepharose 4B beads (17075601, GE Healthcare Life Sciences), for 4-5 hours at 4°C. After washing, beads were further incubated overnight at 4°C with 5-10 mg of *S. pombe* protein extracts prepared in WEB lysis buffer, before analysis by Coomassie blue staining and Western blotting.

### Mass spectrometry and data analysis

Dry TCA precipitates from TAP eluates were denatured, reduced and alkylated. Briefly, each sample was dissolved in 89 μL of TEAB 100 mM. One microliter of DTT 1 M was added and incubation was performed for 30 min at 60°C. A volume of 10 μL of IAA 0.5 M was added (incubation for 30 min in the dark). Enzymatic digestion was performed by addition of 1 μg trypsin (Gold, Promega, Madison USA) in TEAB 100 mM and incubation overnight at 30°C. After completing the digestion step, peptides were purified and concentrated using OMIX Tips C18 reverse-phase resin (Agilent Technologies Inc.) according to the manufacturer’s specifications. Peptides were dehydrated in a vacuum centrifuge.

Samples were resuspended in 9 μL formic acid (0.1%, buffer A) and 2 μL were loaded onto a 15 cm reversed phase column (75 mm inner diameter, Acclaim Pepmap 100® C18, Thermo Fisher Scientific) and separated with an Ultimate 3000 RSLC system (Thermo Fisher Scientific) coupled to a Q Exactive Plus (Thermo Fisher Scientific) via a nano-electrospray source, using a 143-min gradient of 2 to 40% of buffer B (80% ACN, 0.1% formic acid) and a flow rate of 300 nl/min.

MS/MS analyses were performed in a data-dependent mode. Full scans (375 – 1,500 m/z) were acquired in the Orbitrap mass analyzer with a 70,000 resolution at 200 m/z. For the full scans, 3 x 106 ions were accumulated within a maximum injection time of 60 ms and detected in the Orbitrap analyzer. The twelve most intense ions with charge states ≥ 2 were sequentially isolated to a target value of 1 x 105 with a maximum injection time of 45 ms and fragmented by HCD (Higher-energy collisional dissociation) in the collision cell (normalized collision energy of 28%) and detected in the Orbitrap analyzer at 17,500 resolution.

Raw spectra were processed using the MaxQuant environment (v.1.5.5.1) (Cox & Mann, 2008) and Andromeda for database search with label-free quantification (LFQ), match between runs and the iBAQ algorithm enabled (Cox *et al*, 2011). The MS/MS spectra were matched against the UniProt Reference proteome (Proteome ID UP000002485) of *S. pombe* (strain 972 / ATCC 24843) (Fission yeast) (release 2017_10; https://www.uniprot.org/) and 250 frequently observed contaminants as well as reversed sequences of all entries. Enzyme specificity was set to trypsin/P, and the search included cysteine carbamidomethylation as a fixed modification and oxidation of methionine, and acetylation (protein N-term) and/or phosphorylation of Ser, Thr, Tyr residue (STY) as variable modifications. Up to two missed cleavages were allowed for protease digestion. FDR was set at 0.01 for peptides and proteins and the minimal peptide length at 7.

The relative abundance of proteins identified in each affinity purification was calculated as described in (Smits *et al*, 2013). Briefly, label-free quantification (LFQ) intensity based values were transformed to a base 2 logarithmic scale (Log2), to fit the data to a Gaussian distribution and enable the imputation of missing values. Normalized LFQ intensities were compared between replicates, using a 1% permutation-based false discovery rate (FDR) in a two-tailed Student’s t-test (Table S1). The threshold for significance was set to 1 (fold change = 2), based on the FDR and the ratio between TAP and ‘no TAP’ samples. The relative abundance of subunits in each purification eluate was obtained by dividing the LFQ intensity of that interactor (prey) to the LFQ intensity of the TAP purified protein (bait). For scaling purpose (Figure 4B and 6B,C), this ratio was further normalized to that obtained in control conditions and expressed as percentage.

### RNA-seq and data analysis

All strains were done in triplicate. RNA was extracted from 50 mL of exponentially growing cells RNA using TRIzol reagent (15596018, ThermoFisher Scientific). DNA was removed by DNase I digestion, using the TURBO DNA-free™ kit (AM1907, Ambion) and RNA was cleaned using the RNeasy Mini kit (74104, Qiagen). Total RNA quality and concentration was determined using an Agilent Bioanalyzer. Transcripts were purified by polyA-tail selection. Stranded dual-indexed cDNA libraries were constructed using the Illumina TruSeq Stranded mRNA Library Prep kit. Library size distribution and concentration were determined using an Agilent Bioanalyzer. 48 libraries were sequenced in one lane of an Illumina HiSeq 4000, with 1x 50 bp single reads, at Fasteris SA (Plan-les-Ouates, Switzerland). After demultiplexing according to their index barcode, the total number of reads ranged from 6 to 10 million per library.

Adapter sequences were trimmed from reads in the Fastq sequence files. Reads were aligned using HISAT2 (Kim *et al*, 2015), with strand-specific information (--rna-strandness R) and otherwise default options. For all 48 samples, the overall alignment rate was over 95%, including over 90% of reads mapping uniquely to the *S. pombe* genome. Reads were then counted for gene and exon features using htseq-count (Anders *et al*, 2015) in union mode (-- mode union), reverse stranded (--stranded Reverse), and a minimum alignment quality of 10 (--minaqual 10). For all samples, over 95% of reads were assigned to a feature (--type gene). Variance-mean dependence was estimated from count tables and tested for differential expression based on a negative binomial distribution, using DESeq2 (Love *et al*, 2014). Pairwise comparison or one-way analysis of variance were run with a parametric fit and genotype as the source of variation (factor: ‘mutant’ or ‘control’). All computational analyses were run on the Galaxy web platform using the public server at usegalaxy.org (Afgan *et al*, 2018).

## Statistical analysis

Statistical tests were performed using Graphpad Prism. *t*-tests were used when comparing two means. One-way or two-way analyses of variance (ANOVA) were performed for comparing more than two means, across one (for example “genotype”) or two distinct variables (for example “genotype” as a between-subject factors and “time” as a within-subject factor). One-way ANOVAs were followed by Tukey and two-way ANOVAs were followed by Bonferroni post-hoc pairwise comparisons. An α level of 0.01 was used *a priori* for all statistical tests, except otherwise indicated. Comparisons that are statistically significant (*p* ≤ 0.01) are marked with the star sign (*).

## Data availability

The raw sequencing data reported in this paper have been deposited in the NCBI Gene Expression Omnibus under accession number ###.

## Acknowledgments

We thank Kerstin Wagner and Elsa Cesari for invaluable technical assistance and all members of the Helmlinger laboratory for helpful suggestions and discussions. We are grateful to Robin Allshire, Charles Hoffman, Hisao Masukata, Paul Russell, and Fred Winston for kindly sharing strains. A.E.V. is a recipient of a post-doctoral fellowship from the Fondation pour la Recherche Médicale. D.T. is a recipient of a graduate fellowship from the French Ministry for Research and Higher Education (MESR), supported by the Labex EpiGenMed, an « Investissements d’avenir » program (ANR-10-LABX-12-01). This work was supported by funds from the CNRS (ATIP-Avenir), the FP7 Marie Curie Actions (FP7-PEOPLE-2012-CIG/COACTIVATOR), and the Agence Nationale de la Recherche (ANR-15-CE12-0009-01 and ANR-15-CE11-0022-03) to D.H..

