## Supplemental Figures and Tables for "Chaperone-mediated ordered assembly of the SAGA and NuA4 transcription co-activator complexes"

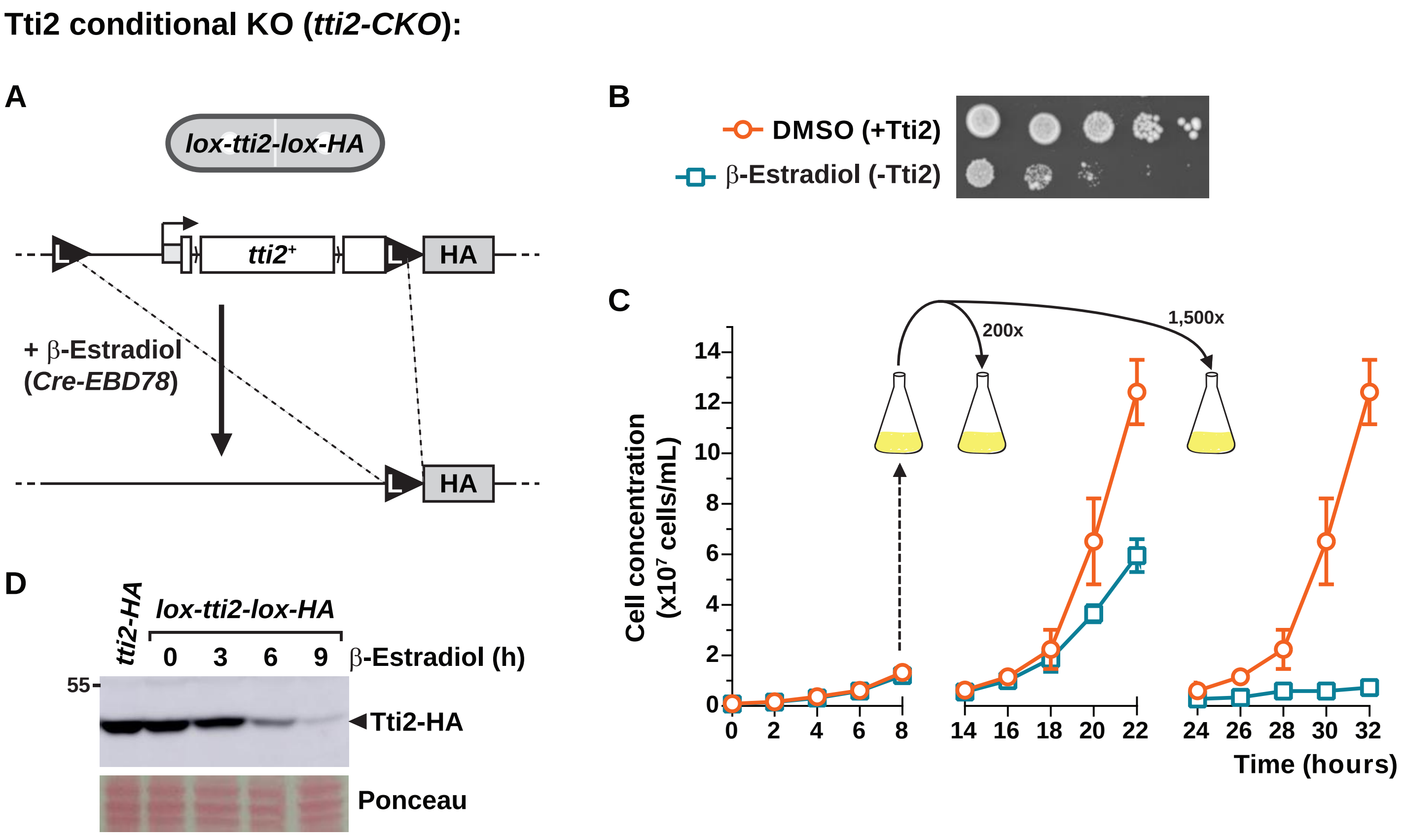


**Supplemental Figure 1. Cre-ER-mediated inducible knock-out of Tti2.**

(A) Illustration of the *tti2*+ locus engineered with loxP sites (L black triangles) flanking the entire gene, before and after β-estradiol-induced Cre recombinase activity.

(B-C) Growth phenotype *tti2-CKO* strains grown in liquid media supplemented with either DMSO (+Tti2, orange circles) or β-estradiol (-Tti2, blue squares). (B) Ten-fold serial dilutions of cells treated for 16 hours with either DMSO or β-estradiol were spotted on rich medium and incubated for 3 days at 32°C. (C) Growth curves from three distinct experiments, in which cells were diluted differently so as to count them during exponential growth, but at different time points in DMSO- or β-estradiol-containing media. Each value represents the average number of cells from three independent replicates with the standard deviation (SD). At least 50 cells from the indicated genotypes were counted at each time point.

(D) Anti-HA Western blot analysis of Tti2-HA expression at different time points upon *tti2*+ knock-out, as compared to a non-loxed tti2-HA strain. Ponceau red staining is used as loading control.

**Tra2 conditional KO (*tra2-CKO*):**

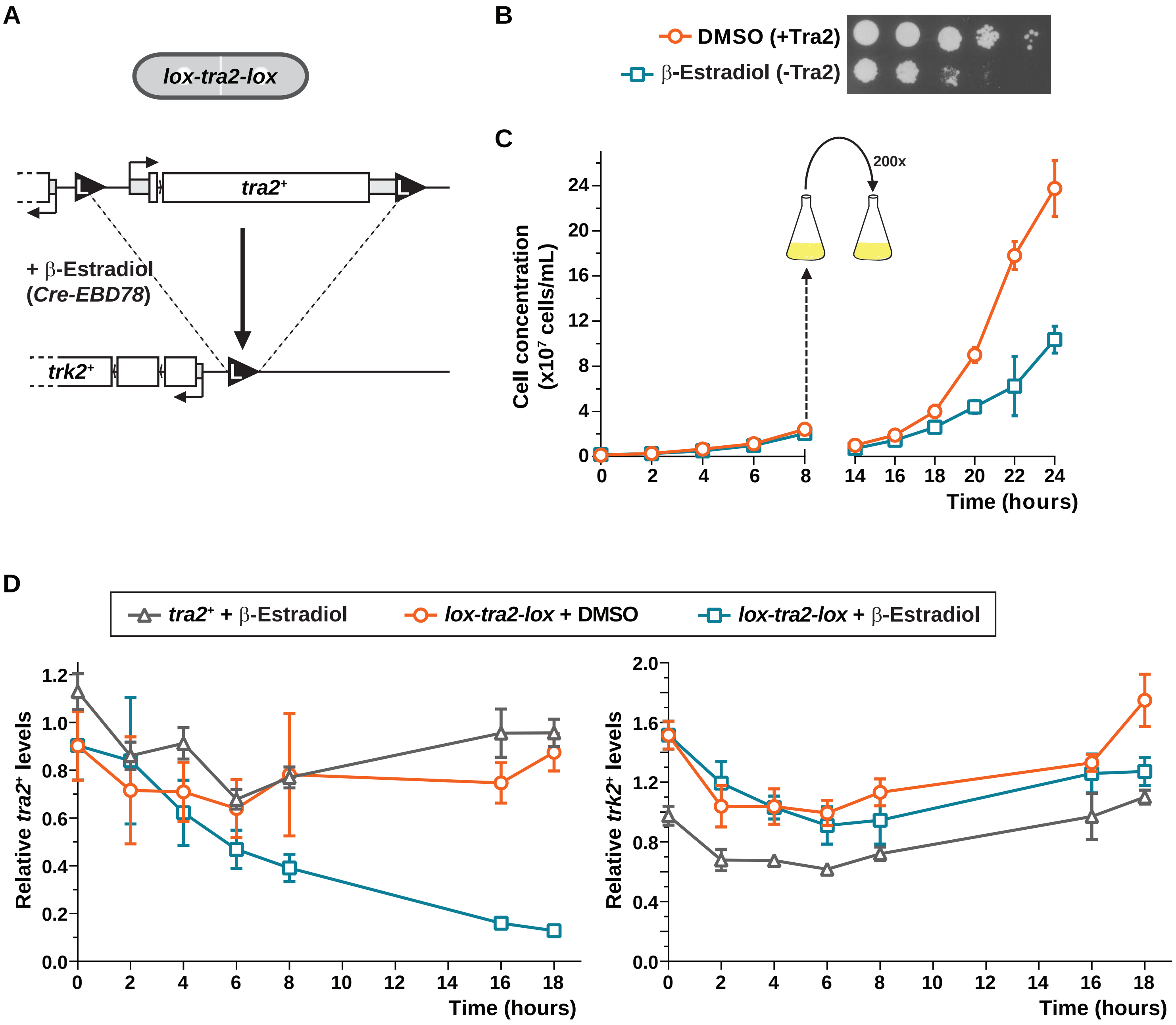


**Supplemental Figure 2. Cre-ER-mediated inducible knock-out of Tra2.**

(A) Illustration of the *tra2*+ locus engineered with loxP sites (L black triangles) flanking the entire gene, before and after β-estradiol-induced Cre recombinase activity.

(B-C) Growth phenotype *tra2-CKO* strains grown in liquid media supplemented with either DMSO (+Tra2, orange circles) or β-estradiol (-Tra2, blue squares). (B) Ten-fold serial dilutions of cells treated for 18 hours with either DMSO or β-estradiol were spotted on rich medium and incubated for 3 days at 32°C. (C) Growth curves from two distinct experiments, in which cells were diluted differently so as to count them during exponential growth, but at different time points in DMSO- or β-estradiol-containing media. Each value represents the average number of cells from three independent replicates with the standard deviation (SD). At least 50 cells from the indicated genotypes were counted at each time point.

(D) RT-qPCR analysis of the expression of *tra2+* (left) and of its divergently transcribed 5' neighbouring gene, *trk2+* (right), at different time points upon tra2+ knock-out. A wild-type control strain grown in β-estradiol (grey triangles) and a *tra2-CKO* strain grown either in DMSO (orange circles) or in β-estradiol (blue squares) were analyzed. Each value represents mean mRNA levels from three independent RT-qPCR experiments with the SD. *act1+* served as a control for normalization across samples. Values from one control experiment were set at 1 to allow comparisons across culture conditions and mutant strains.

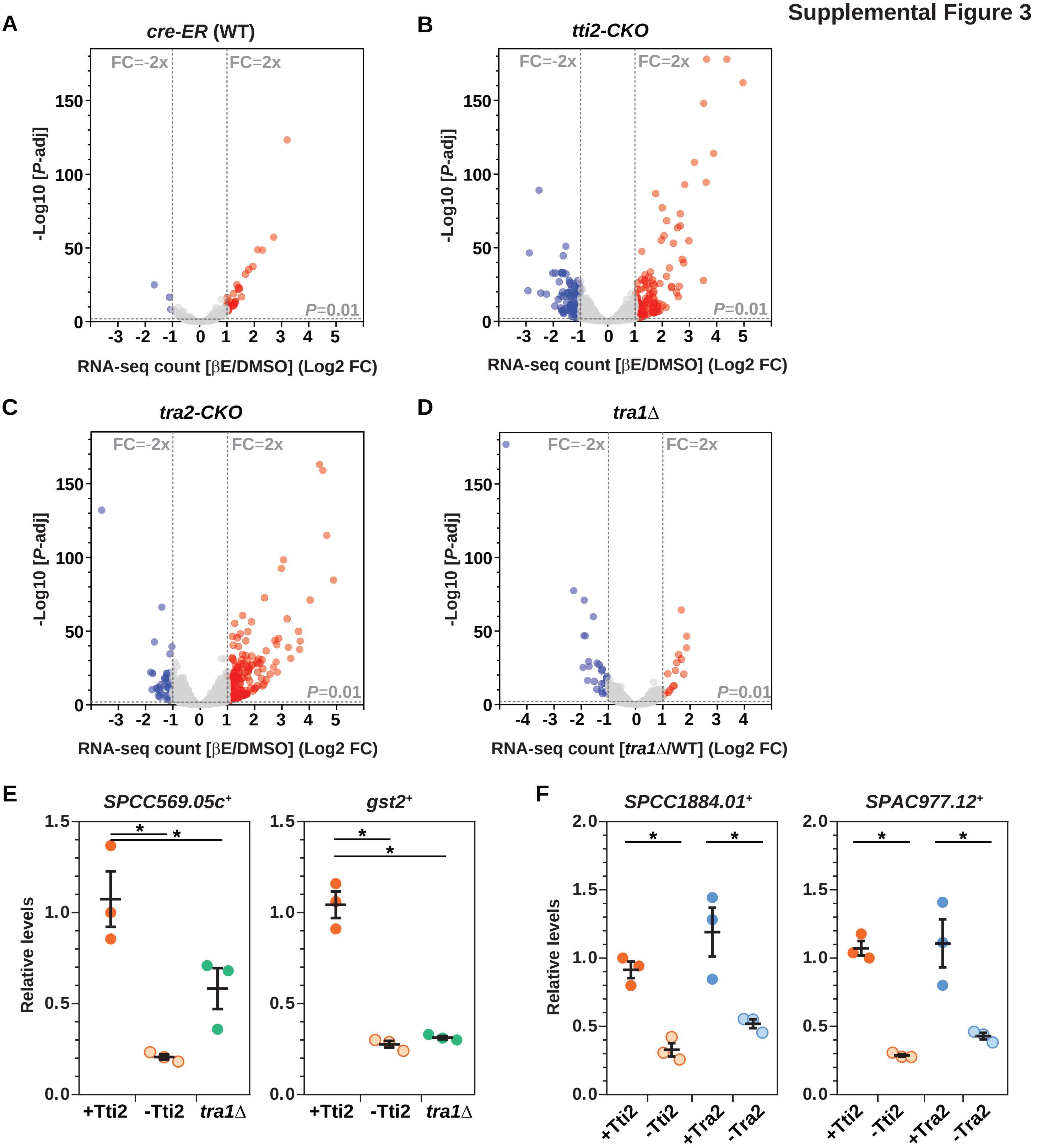


**Supplemental Figure 3. Transcriptome analyses of *tti2-CKO*, *tra2-CKO* and *tra1*Δ mutants.**
(A-D) Volcano plots of RNA-seq data comparing the fold change versus the adjusted *P* value, calculated using the DESeq2 package. RNA-seq were performed in triplicate from control *cre-ER* (WT) (A), inducible *tti2+* knock-out (*tti2-CKO*) (B), inducible *tra2+* knock-out (*tra2-CKO*) (C), and *tra1*Δ mutants (D). Differential gene expression analysis was performed comparing cells treated with either DMSO or β-estradiol (βE) for the *cre-ER*, *tti2-CKO*, and *tra2-CKO* strains. *tra1*Δ mutants were compared to isogenic wild-type (WT) control cells. Up- and down-regulated genes with a fold change ≥ 2 and a *P* value ≤ 0.01 are coloured in red and blue, respectively, and these thresholds are shown as grey dashed lines.
(E-F) Expression of Tra1- and Tra2-dependent genes upon conditional knock-out of *tti2+*, *tra2+*, or in *tra1*Δ mutants. mRNA levels of *SPCC569.05c+*, *gst2+*, *SPCC1884.01+*, and *SPAC977.12+* were measured using RT-qPCR of RNA extracted from *tti2-CKO*, *tra2-CKO*, and *tra1*Δ strains grown to exponential phase in rich medium supplemented with DMSO (+Tti2 or +Tra2) or β-estradiol (-Tti2 or -Tra2). Each value represents mean mRNA levels from three independent experiments with the SEM. *act1+* served as a control for normalization across samples.
Values from one control experiment were set at 1 to allow comparisons across culture conditions and mutant strains. Statistical significance was determined by one-way ANOVA followed by Tukey's multiple comparison tests (*$P < 0.05$).

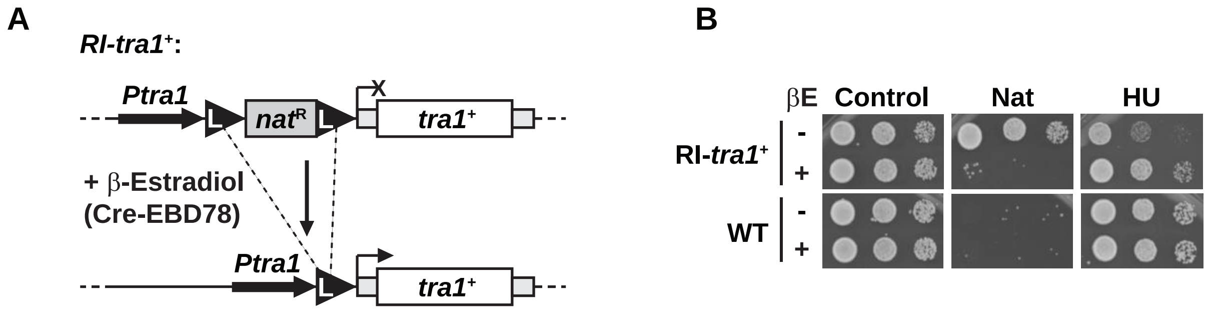


**Supplemental Figure 4. Genetic assay to follow the fate of Tra1 upon *de novo* synthesis.**

(A) Illustration of the strategy to follow the fate of newly synthesized Tra1 (Recombination Induced-*tra1*+). A strong transcription terminator sequence from the *Ashbya gossypii TEF1* gene, included in the antibiotic resistance cassette, flanked by two *loxP* sites (L black triangles) was inserted between *tra1*+ promoter and ORF. Addition of β-estradiol allows recombination-induced *de novo* expression of Tra1 at endogenous levels.

(B) Growth phenotype of control WT and *RI-tra1*+ strains, grown in liquid media and treated with either DMSO (-) or β-estradiol (+). Ten-fold serial dilutions were spotted on rich medium (control), supplemented with Nourseothricin (Nat) or 10 mM hydroxyurea (HU), and incubated for 3 days at 32°C.
.

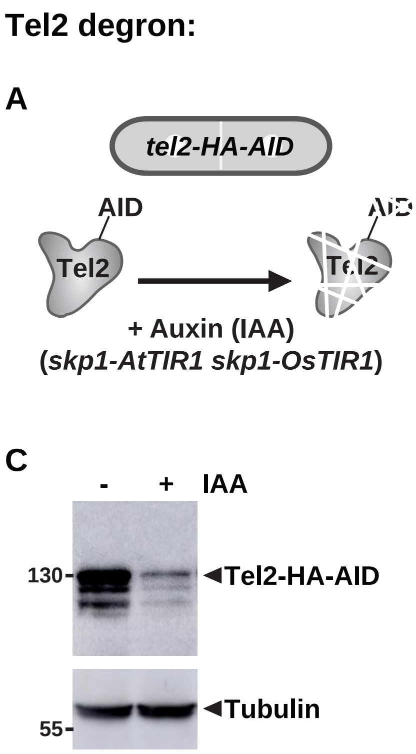


**Supplemental Figure 5. Auxin-mediated inducible depletion of Tel2.**

(A) Illustration of the auxin-inducible degron (AID) system for endogenous Tel2 depletion, using strains in which *S. pombe* Skp1 is fused to *Arabidopsis thaliana* Tir1 and *Oryza sativa* Tir1. Addition of the plant hormone auxin to the media (IAA; indolacetic acid) allows the rapid and conditional degradation of Tel2.

(B) Growth phenotype cells depleted for Tel2. *tel2-HA-AID skp1-AtTIR1 skp1-OsTIR1* cells were grown to exponential phase in minimal medium supplemented with either ethanol (EtOH; +Tel2) or auxin (IAA; -Tel2). Ten-fold serial dilutions were then spotted on corresponding solid media and grown for 3 days at 25°C. WT and *skp1-AtTIR1 skp1-OsTIR1* strains were used as controls.

(C) Anti-HA Western blot analysis of Tel2-HA-AID expression 16 hours after adding auxin. An anti-tubulin antibody served as loading control.

### Supplemental Figure 6

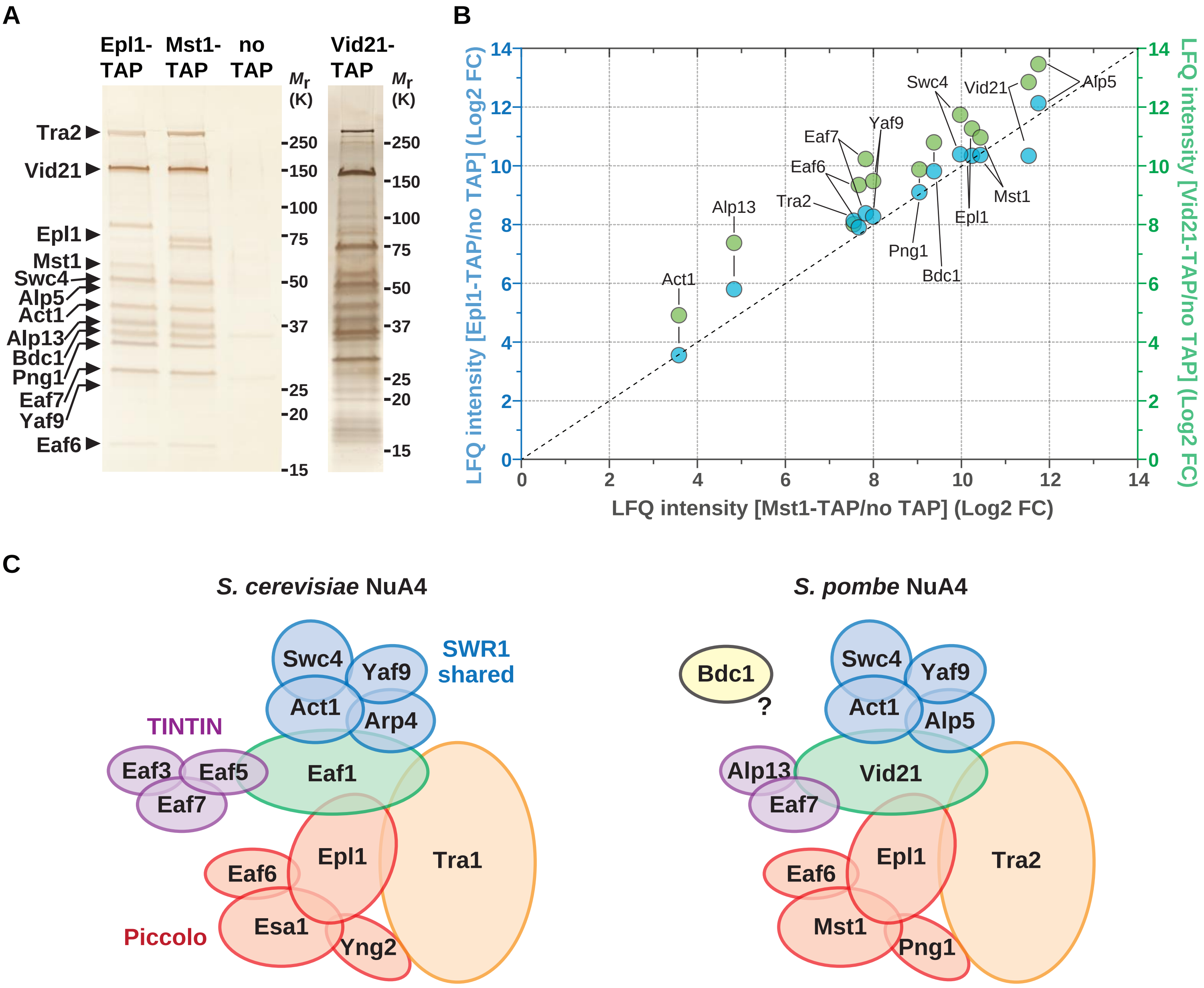


**Supplemental Figure 6. Characterization of the *S. pombe* NuA4 complex.**

(A) Silver staining analysis of NuA4 complexes purified using Mst1, Epl1, or Vid21 as baits. A non-tagged strain (no TAP) was used as a control for background. The position of each NuA4 subunit is annotated on the gel according to their predicted molecular weight.

(B) Tandem mass spectrometry analyses (LC-MS/MS) of tandem affinity purified Mst1, Epl1, and Vid21 from (A). Each individual point represents the LFQ intensity ratio of a protein in TAP eluates relative to a ‘no TAP’ control. Blue compares Epl1 with Mst1 eluates, whereas green compares Vid21 with Mst1 eluates. The black dashed line represents a 1:1 ratio.

(C) Comparison of NuA4 complex subunit composition between *S. cerevisiae* and *S. pombe*.

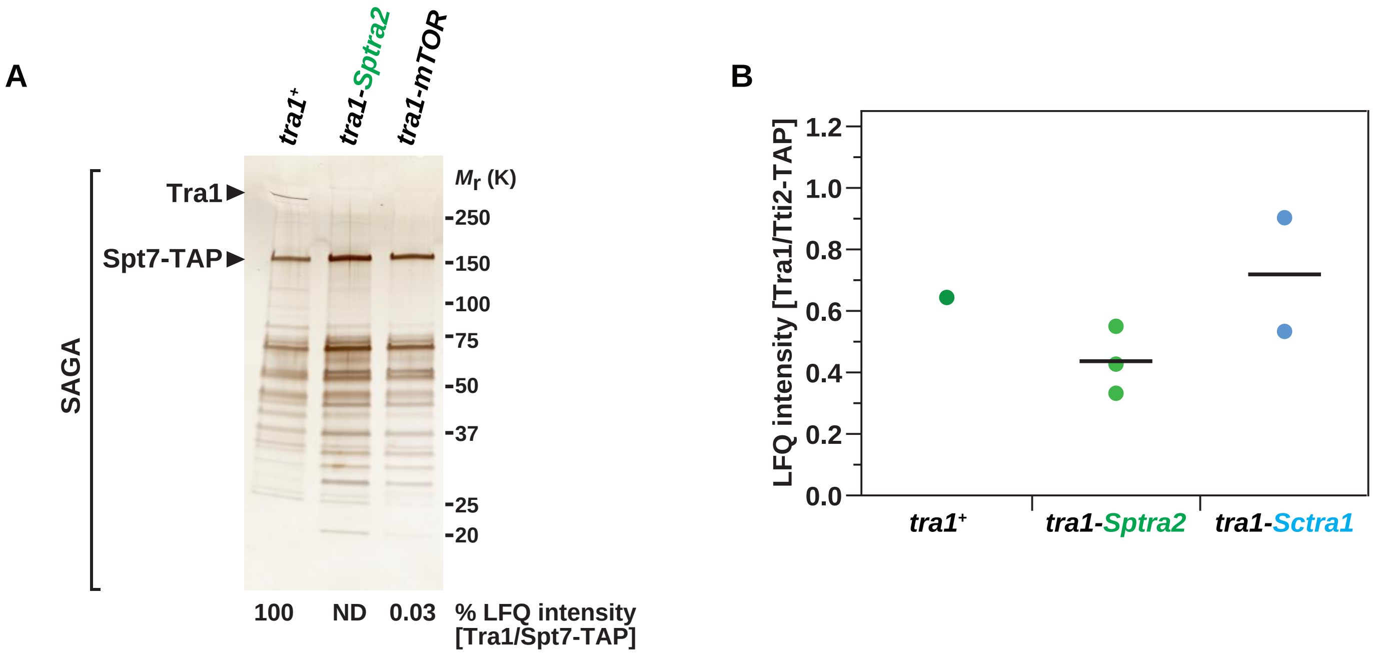


**Supplemental Figure 7. The CSI region of Tra1 is required for interaction with SAGA, but not with TTT.**

(A) Residues 2623-2676 from *S. pombe* Tra1 were swapped with the homologous region from either *S. pombe* Tra2 (residues 2564-2617, green), to create the *tra1-Sptra2* allele, or *Homo sapiens* mTOR (residues 1456-1504), to create the *tra1-mTOR* allele. Silver staining analysis of SAGA complexes purified from WT, *tra1-Sptra2*, and *tra1-mTOR* strains, using Spt7 as the bait. Numbers at the bottom of the gel represent LFQ intensity ratios of Tra1 to Spt7, from LC-MS/MS analyses of purified SAGA complexes. Values for each mutant are expressed as percentage of WT SAGA (ND: not detected). Shown are gels that are representative of two independent experiments.

(B) LC-MS/MS analysis of TTT complexes purified from WT, *tra1-Sptra2*, and *tra1-Sctra1* strains (see Figure 5B), using Tti2 as the bait. LFQ intensity ratios of Tra1 to Tti2 from 1-3 biological replicates are plotted individually with the mean (black bar).

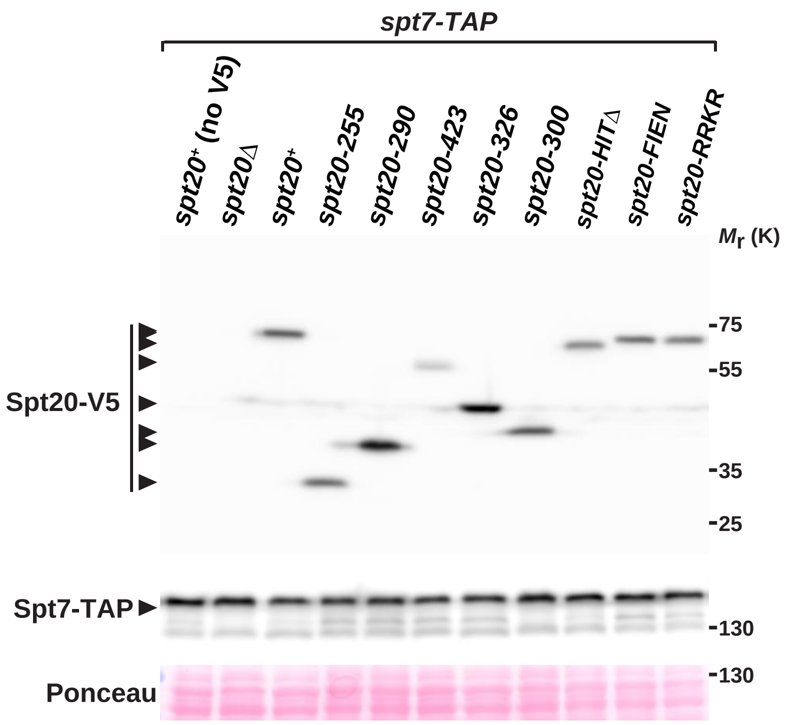


**Supplemental Figure 8. Western blot controls for SAGA TAPs in Spt20 truncation and point mutants.**

Anti-V5 and anti-HA Western blot analyses of Spt20-V5 (upper blot) and Spt7-TAP (lower blot) in a fraction of the input used for TAPs shown in Figure 5. Ponceau red staining is used as loading control.

**Supplemental Table S1.** Summary of proteins identified by tandem affinity purifications of Tti2. Shown are results of LC-MS/MS analyses of total protein mixtures from purification eluates. Numbers indicate the LFQ intensity fold change (FC) of Tti2-TAP to 'no TAP' control, averaged from four independent replicates. Shown are proteins enriched in Tti2 purifications at least 2-fold, with a $q < 0.05$.

| Name | mean FC (Log2) | -Log10[q value] |
|---|---:|---:|
| Tti2 (bait) | 10.73123975 | 2.394611602 |
| Tti1 | 8.65016021 | 2.394611602 |
| Tra1 | 8.606427752 | 2.33510836 |
| Tor2 | 7.705570879 | 2.394611602 |
| Tel2 | 7.347468728 | 2.33510836 |
| Asa1 | 6.638643717 | 1.807795544 |
| Tra2 | 6.496988347 | 1.468259394 |
| Wat1 | 4.884663889 | 1.380746842 |
| Tor1 | 4.533265776 | 1.468259394 |
| Rad3 | 3.974289999 | 1.380746842 |
| Rvb2 | 1.561711276 | 2.33510836 |

**Supplemental Table S2: List of *S. pombe* and *S. cerevisiae* strains used in this study.**

| Strain | Genotype | | Source |
|---|---|---|---|
| DHP42 | *h+* | | Lab stock |
| DHP43 | *h-* | | Lab stock |
| DHP1139 | *h-* | *tti2-GLY6-HA3-TAP2::kanMX6* | This study |
| DHP609 | *h-* | *tra1Δ::kanMX6* | This study |
| DHP1235 | *h-* | *leu1-32 ars1::pRad15-Cre-EBD-LEU2* | Derived from A9823 (R. Allshire) |
| DHP1487 | *h-* | *leu1-32 ars1::pRad15-Cre-EBD-LEU2 lox::kanMX6::tti2-lox-HA* | This study |
| DHP1488 | *h-* | *leu1-32 ars1::pRad15-Cre-EBD-LEU2 lox::kanMX6::tra2::lox* | This study |
| DHP1489 | *h-* | *leu1-32 ars1::pRad15-Cre-EBD-LEU2 lox::kanMX6::tti2-lox-HA spt7-MYC13::natMX6* | This study |
| DHP1490 | *h-* | *leu1-32 ars1::pRad15-Cre-EBD-LEU2 lox::kanMX6::tti2-lox-HA epl1-MYC13::natMX6* | This study |
| DHP782 | *h-* | *spt7-HA3-TAP2::kanMX6* | Lab stock |
| DHP1293 | *h-* | *epl1-HA3-TAP2::kanMX6* | This study |
| DHP1295 | *h-* | *mst1-HA3-TAP2::kanMX6* | This study |
| DHP1492 | *h-* | *tra1Δ::hphMX6 spt7-HA3-TAP2:kanMX6* | This study |
| DHP1491 | *h-* | *leu1-32 ars1::pRad15-Cre-EBD-LEU2 lox::kanMX6::tti2-lox-HA spt7-HA3-TAP2::kanMX6* | This study |
| DHP1493 | *h-* | *leu1-32 ars1::pRad15-Cre-EBD-LEU2 lox::kanMX6::tti2-lox-HA mst1-HA3-TAP2::kanMX6* | This study |
| DHP1494 | *h-* | *leu1-32 ars1::pRad15-Cre-EBD-LEU2 lox::kanMX6::tra2::lox epl1-HA3-TAP2::kanMX6* | This study |
| DHP1495 | *h-* | *leu1-32 ars1::pRad15-Cre-EBD-LEU2 lox::kanMX6::tra2::lox vid21-TAP2::natMX6* | This study |
| DHP1188 | *h-* | *ade6::ade6-Padh15-skp1-OsTIR1-natMX6-Padh15-skp1-AtTIR1-2NLS* | Derived from HM2985 (H. Masukata) |
| DHP1179 | *h-* | *ade6::ade6-Padh15-skp1-OsTIR1-natMX6-Padh15-skp1-AtTIR1-2NLS tel2-HA3-AID3::kanMX6* | This study |
| DHP1496 | *h-* | *leu1-32 ars1::pRad15-Cre-EBD-LEU2 natMX6::RI-tra1 spt7-HA3-TAP2::kanMX6 ade6::ade6-Padh15-skp1-OsTIR1-natMX6-Padh15-skp1-AtTIR1-2NLS tel2-HA3-mAID3::kanMX6* | This study |
| DHP1497 | *h+* | *leu1-32 ars1::pRad15-Cre-EBD-LEU2 natMX6::RI-tra1 spt7-HA3-TAP2::kanMX6 hsp90-201::kanMX6* | This study |
| DHP1498 | *h-* | *leu1-32 ars1::pRad15-Cre-EBD-LEU2 natMX6::RI-tra1 spt7-HA3-TAP2::kanMX6* | This study |
| DHP852 | *h-* | *ura4::fbp1-lacZ hsp90-201::kanMX6* | Derived from CHP879 (C. Hoffman) |
| DHP1521 | *h+* | *hsp90-26* | Derived from RA1193 (P. Russell) |
| DHP1522 | *h+* | *hsp90-26 spt7-HA3-TAP2::kanMX6* | This study |
| DHP1432 | *h-* | *tra1-Sptra2* | This study |
| DHP1425 | *h-* | *tra1-Sctra1* | This study |
| DHP1483 | *h-* | *FLAG3-tra1-Sptra2* | This study |
| DHP1532 | *h-* | *FLAG3-tra1-Sctra1* | This study |
| DHP1523 | *h-* | *FLAG3-tra1 spt7-HA3-TAP2::kanMX6* | This study |
| DHP1514 | *h-* | *FLAG3-tra1-Sptra2 spt7-HA3-TAP2::kanMX6* | This study |
| DHP1515 | *h-* | *FLAG3-tra1-Sctra1 spt7-HA3-TAP2::kanMX6* | This study |
| DHP1373 | *h+* | *ura4-D18 leu1-32 ade6-M216 spt7-HA3-TAP2::kanMX6 tra1-mTOR* | This study |
| DHP1434 | *h-* | *tra1-Sptra2 tti2-GLY6-HA3-TAP2::kanMX6* | This study |
| DHP1424 | *h-* | *tra1-Sctra1 tti2-GLY6-HA3-TAP2::kanMX6* | This study |

| | | | |
|---|---|---|---|
| DHP1318 | *h-* | *spt20Δ::natMX6 spt7-HA3-TAP2::kanMX6* | This study |
| DHP1519 | *h-* | *spt20-PK3::hphMX6* | This study |
| DHP1358 | *h-* | *spt20-255-PK3::hphMX6 spt7-HA3-TAP2::kanMX6* | This study |
| DHP1365 | *h-* | *spt20-300-PK3::hphMX6 spt7-HA3-TAP2::kanMX6* | This study |
| DHP1380 | *h-* | *spt20-290-PK3::hphMX6 spt7-HA3-TAP2::kanMX6* | This study |
| DHP1382 | *h-* | *spt20-423-PK3::hphMX6 spt7-HA3-TAP2::kanMX6* | This study |
| DHP1401 | *h-* | *spt20-326-PK3::hphMX6 spt7-HA3-TAP2::kanMX6* | This study |
| DHP1409 | *h-* | *ura4-D18 ade6-M216 spt20-PK3::hphMX6 spt7-HA3-TAP2::kanMX6* | This study |
| DHP1421 | *h-* | *ura4-D18 ade6-M216 spt20-HITΔ-PK3::hphMX6 spt7-HA3-TAP2::kanMX6* | This study |
| DHP1444 | *h-* | *ura4-D18 ade6-M216 spt20-RRKR-PK3::hphMX6 spt7-HA3-TAP2::kanMX6* | This study |
| DHP1446 | *h-* | *ura4-D18 ade6-M216 spt20-FIEN-PK3::hphMX6 spt7-HA3-TAP2::kanMX6* | This study |
| DHP1516 | *h-* | *spt20Δ::kanMX6* | This study |
| DHP1517 | *h-* | *spt20-HITΔ-PK3::hphMX6* | This study |
| DHP1448 | *MATa* | *ura3D0 leu2D1 trp1D63 hisD4-917 lys2-173R2 HA-SPT7-TAP::TRP1* | FY2031 (F. Winston) |
| DHP1460 | *MATa* | *ura3D0 leu2D1 trp1D63 hisD4-917 lys2-173R2 HA-SPT7-TAP::TRP1 spt20-D492aa-PK3::hphMX6* | This study |
| DHP1462 | *MATa* | *ura3D0 leu2D1 trp1D63 hisD4-917 lys2-173R2 HA-SPT7-TAP::TRP1 spt20-D380aa-PK3::hphMX6* | This study |
| DHP1524 | *MATa* | *ura3D0 leu2D1 trp1D63 hisD4-917 lys2-173R2 HA-SPT7-TAP::TRP1 spt20-D474aa-PK3::hphMX6* | This study |
| DHP1520 | *h-* | *ura4-D18 tra1Δ::ura4 spt7-HA3-TAP2::kanMX6 sgf11-MYC13::natMX6* | This study |
| DHP1499 | *h-* | *leu1-32 ars1::pRad15-Cre-EBD-LEU2 natMX6::RI-tra1 spt7-HA3-TAP2::kanMX6 sgf11-MYC13::natMX6* | This study |
| DHP2 | *h-* | *ura4-D18 ade6-M210 ada1-HA3-TAP2::kanMX6* | Lab stock |
| DHP486 | *h-* | *ura4-D18 ade6-M216 sgf73Δ::ura4 ada1-HA3-TAP2::kanMX6* | This study |

**Supplemental Table S3. List of oligonucleotides used in this study.**

| Gene | Name | Description[a] | Coordinates[b] | Sequence |
|---|---|---|---|---|
| | DHO 1176 | Subcloning HA tag (DHB137) | | AGCTGAAGCTTCGTACGCTGCAGGTACCCATACGATGTTCCTGACTATGC |
| | DHO 1177 | Subcloning HA tag (DHB137) | | GCACCAGCACCAGCACCGGCACCGGCAGCGTAATCTGGAACGTCATATGG |
| | DHO 1513 | Subcloning V5-PK tag (DHB123) | | TCGATGATCCCAAACCCATTGTTGGGTTTGGACGGT |
| | DHO 1514 | Subcloning V5-PK tag (DHB123) | | TCGAACCGTCCAAACCCAACAATGGGTTTGGGATCA |
| *act1* | DHO 90 | RT-qPCR | +10 to +29 | GAAATCGCAGCGTTGGTTAT |
| *act1* | DHO 91 | RT-qPCR | +186 to +167 | ACGCTTGCTTTGAGCTTCAT |
| *epl1* | DHO 1695 | C-terminal tagging / pFA6a | | CAATTATACTCGATGATGAATCTGATGCTTCTATAATGTTTCGAGCTAGCTTGCTAAACGATGATATGGGTACTCAGTCACGGATCCCCGGGTTAATTAA |
| *epl1* | DHO 1696 | C-terminal tagging / pFA6a | | CGAGGAGAAATAGTTAAAATAAAATTTGATCGAAATCTTTAGTAATGGAAATATTATAAATACCATAGATAAAAAGCGTTGAATTCGAGCTCGTTTAAAC |
| *gst2* | DHO 1900 | RT-qPCR | +233 to +252 | TGGCTGACAAGTATGATACG |
| *gst2* | DHO 1901 | RT-qPCR | +400 to +382 | TGACTGCAGATACCACTGG |
| *mei2* | DHO 188 | ChIP-qPCR | -1946 to -1924 | GAGCTTACTGACATCGCATCA |
| *mei2* | DHO 189 | ChIP-qPCR | -1721 to -1701 | TCAACAGCAGCTTTCAGTGG |
| *mst1* | DHO 1699 | C-terminal tagging / pFA6a | | AAAGAAGAAGGATCAATGGAGATTTGTTGGCAGATTGGCAACCTCCTGTTTTTCACCCTTCCCAACTACGATTTGGATGGCGGATCCCCGGGTTAATTAA |
| *mst1* | DHO 1700 | C-terminal tagging / pFA6a | | ACATGTGGGCTAGTATACGATATCCAGCTATCCTGCAAAATGGAGCAGCCGAAAAAACCAAGGCAATAGCAGAGGCTTTTGAATTCGAGCTCGTTTAAAC |
| *pho84* | DHO 1300 | ChIP-qPCR | -477 to -458 | CCGCGTTACCTTAATCCTGA |
| *pho84* | DHO 1301 | ChIP-qPCR | -250 to -269 | AGCTTTGGTGTTGACGTTCC |
| *sgf11* | DHO 1715 | C-terminal tagging / pFA6a | | AGAAATGTAAAGGCAGGTCACTTTCAAACGCTACCTCGTACTCAACTCTTTTCGAAGATGACAATGCAGACGAAGAGGACCGGATCCCCGGGTTAATTAA |
| *sgf11* | DHO 1716 | C-terminal tagging / pFA6a | | ATAATATTGTAGTTCGTCTTACTGGGAAAGACATACATGTGCATAATGAGAATAAACCTTAAAACAAGCAACAAATAAAAGAATTCGAGCTCGTTTAAAC |
| *SPAC977.12* | DHO 1890 | RT-qPCR | +123 to +142 | TGCAATGGGTGGTACAATCG |
| *SPAC977.12* | DHO 1891 | RT-qPCR | +311 to +292 | AGTTTTAGGACATCAGCAGG |
| *SPCC1884.01* | DHO 1892 | RT-qPCR | +929 to +949 | AGAACATCATCGTTCGCTTGG |
| *SPCC1884.01* | DHO 1893 | RT-qPCR | +1111 to +1093 | TTGGATATCGACCTCTTTGG |
| *SPCC569.05c* | DHO 1896 | RT-qPCR | +124 to +143 | TCACAGCTTTCTCATTCTGC |
| *SPCC569.05c* | DHO 1897 | RT-qPCR | +343 to +324 | AGACAAACCGATCAGGATCC |
| *spt20* | DHO 849 | Deletion / pFA6a | | TTAACCAGCTTATAATTTTAATTGTAAACATAGGGGGAATATTGTTTTACAATTACTGGTATAGTTCGTTGAACTAAGAGCGGATCCCCGGGTTAATTAA |
| *spt20* | DHO 850 | Deletion & C-terminal tagging / pFA6a | | TATGCAACAAAAAAAATAAAATTTTAAAAATAATAAATAAAAAACTAAATTCGTGTATTAAATGATGAAGCATTTGAGAGGAATTCGAGCTCGTTTAAAC |
| *spt20* | DHO 1592 | C-terminal tagging / pFA6a | | TGACGGTTGCACGAAATCTTCATTTTCAACAACTCCAAATGAGACAGCAGCAACAACAGCAGCAAATGAAGGCTGAAAGGCGGATCCCCGGGTTAATTAA |
| *spt20* | DHO 1558 | C-terminal deletion (256-474) / pFA6a | +764 to +843 | AGTCTAAAGCGGAAATGATTCAATTTATGAACCAGTTGGCCGACTCTGCACCACCATGCACCAGAAAGAAACCTCAAGGACGGATCCCCGGGTTAATTAA |
| *spt20* | DHO 1559 | C-terminal deletion (256-474) / pFA6a | +923 to +844 | CGTTTTCTTTGGTCATCCATTAGTAACAAAAGGTTCTCTTTTTCAAGTCTTTCTGCTTCTTCATCAGCAAGTTGGGCTGAGAATTCGAGCTCGTTTAAAC |
| *spt20* | DHO 1560 | C-terminal deletion (291-474) / pFA6a | +769 to +948 | CAGAAAGACTTGAAAAAGAGAACCTTTTGTTACTAATGGATGACCAAAGAAAACGCGATTTTCAACCTACCTTTCAAAGACGGATCCCCGGGTTAATTAA |
| *spt20* | DHO 1561 | C-terminal deletion (291-474) / pFA6a | +1028 to +949 | GCTTTTTGTTGCTGTTGCATTTGCATTTGCCTTTGTTGTAGAATAGCTCGTTTACGCCGCACATTCTCAATAAATTGTAGGAATTCGAGCTCGTTTAAAC |
| *spt20* | DHO 1650 | C-terminal deletion (301-474) / pFA6a | +902 to +981 | TAATGGATGACCAAAGAAAACGCGATTTTCAACCTACCTTTCAAAGACTACAATTTATTGAGAATGTGCGGCGTAAACGACGGATCCCCGGGTTAATTAA |
| *spt20* | DHO 1651 | C-terminal deletion (301-474) / pFA6a | +1061 to +982 | GGTTGCTGAGCTTTCGGTGATTGCTGCTGCTGCGCTTTTTGTTGCTGTTGCATTTGCATTTGCCTTTGTTGTAGAATAGCGAATTCGAGCTCGTTTAAAC |
| *spt20* | DHO 1652 | C-terminal deletion (327-474) / pFA6a | +980 to +1059 | GAGCTATTCTACAACAAAGGCAAATGCAAATGCAACAGCAACAAAAAGCGCAGCAGCAGCAATCACCGAAAGCTCAGCAACGGATCCCCGGGTTAATTAA |
| *spt20* | DHO 1653 | C-terminal deletion (327-474) / pFA6a | +1139 to +1060 | ATACTACTAGGTGGTAGACGTTGGATTTTGGGTGTAGTTTTTCTTTGAACTGGAGCAGACTGAACCAAATGTGCGGGTGGGAATTCGAGCTCGTTTAAAC |
| *spt20* | DHO 1562 | C-terminal deletion (424-474) / pFA6a | +1355 to +1434 | GAGAATCCGTGGATTTGATAAAAATACGACAACTTGCAATTTTGTTTCAACAGCGTGCCTCTCAATTAAAAGCTAGAGGTCGGATCCCCGGGTTAATTAA |
| *spt20* | DHO 1563 | C-terminal deletion (424-474) / pFA6a | +1514 to +1435 | ACCGTCATTACTGTTGCCAAATCAGTACCAGCGGCTATTGCTTGACGATTCAGAATTTCAGTAATTTGTTCCCGTGTAGCGAATTCGAGCTCGTTTAAAC |
| *spt20* | DHO 1485 | N-terminal deletion (1-241) / nmt-HA | +805 to +884 | TTTTCAAGTCTTTCTGCTTCTTCATCAGCAAGTTGGGCTGATCCTTGAGGTTTCTTTCTGGTGCATGGTGGTGCAGAGTCGCACTGAGCAGCGTAATCTG |
| *spt20* | DHO 1486 | N-terminal deletion (1-241) / pFA6a | -80 to -1 | TTAACCAGCTTATAATTTTAATTGTAAACATAGGGGGAATATTGTTTTACAATTACTGGTATAGTTCGTTGAACTAAGAGGAATTCGAGCTCGTTTAAAC |
| *spt20* | DHO 1672 | Internal deletion (290-300) / pKS-ura4 | +769 to +948 | CAGAAAGACTTGAAAAAGAGAACCTTTTGTTACTAATGGATGACCAAAGAAAACGCGATTTTCAACCTACCTTTCAAAGACGCCAGGGTTTTCCCAGTCACGAC |
| *spt20* | DHO 1673 | Internal deletion (290-300) / pKS-ura4 | +1061 to + 982 | GGTTGCTGAGCTTTCGGTGATTGCTGCTGCTGCGCTTTTTGTTGCTGTTGCATTTGCATTTGCCTTTGTTGTAGAATAGCAGCGGATAACAATTTCACACAGGA |
| *spt20* | DHO 1660 | RIP-qPCR | +1425 to +1444 | AAGCTAGAGGTGCTACACGG |
| *spt20* | DHO 1661 | RIP-qPCR | +1563 to +1543 | TGTTGCTGCTGTCTCATTTGG |
| | DHO 1759 | Subcloning HIT-BamHI (DHB179 & DHB193) | | ATCGGATCTGGTTCCGCGTG |
| | DHO 1760 | Subcloning HIT-XhoI (DHB179 & DHB193) | | CAGTCACGATGCGGCCGCT |
| *SPT20* | DHO 1845 | C-terminal tagging / pFA6a | | GTACTTTAATACAATATATATATATATATATATATATATATATATATATATATAAGGAATGATAACTCTATTTAAGTAGAGAATTCGAGCTCGTTTAAAC |
| *SPT20* | DHO 1847 | C-terminal deletion (380-604) / pFA6a | +1061 to +1140 | AAAAATCCTTCATTCACGAACATCGTGCTGAGTCTACGAGAGAAGGCACGAAGGGCGTTGTAGGGCATATCGAAGAACGCCGGATCCCCGGGTTAATTAA |
| *SPT20* | DHO 1849 | C-terminal deletion (492-604) / pFA6a | +1397 to 1476 | CTACCATGGGTAACGATAATAATCAATTTAGCAGGCTGAAATTTATTGAACAATGGAGAATAAATAAAGAAAAAAGAAAACGGATCCCCGGGTTAATTAA |
| *SPT20* | DHO 1964 | C-terminal deletion (474-604) / pFA6a | +1343 to + 1422 | CTAATGGCAATCAAGTAGCTTTGGCAGCGGCAGCGGCTGCGGCTGCAGTGGGATCTACCATGGGTAACGATAATAATCAACGGATCCCCGGGTTAATTAA |
| *ssa2* | DHO 1242 | ChIP-qPCR | -429 to -410 | CACGCTCTTCCAGGACTTTC |
| *ssa2* | DHO 1243 | ChIP-qPCR | -248 to -267 | TCTCGAAGTTTCCCAAAACC |
| *tra1* | DHO 1424 | Internal deletion (2624-2674) / pKS-ura4 | +8022 to +8101 | GATTCACGATTATTAGACCAATCGCTAATTCTCCAAGCACACTCGAGCAATAGTTCAGAAGATCCCTCGTGCTTCGCCAAAGCGGATAACAATTTCACACAGGA |
| *tra1* | DHO 1425 | Internal deletion (2624-2674) / pKS-ura4 | +7790 to +7869 | AAGATGATATGTTTTATGGCCATTGGCGACGAAACTGTAAGTACTTGCAAACTCAAGTTGCTCTCTCGTATGAGCAACTTCGCCAGGGTTTTCCCAGTCACGAC |
| *tra1* | DHO 1764 | CSI sgRNA-1 | +7935 to +7916 | ATAGTTGCTGTTGCCAAAAAACATAACCTGTACCGAAGAAAATGGCTTCACTGCGAGCCTgttttagagctagaaatagcaag |
| *tra1* | DHO 1763 | CSI sgRNA-2 | +7862 to +7881 | ATAGTTGCTGTTGCCAAAAAACATAACCTGTACCGAAGAAAGCAACTTGGAATGTGGGGAgttttagagctagaaatagcaag |
| *tra1* | DHO 1819 | ATG sgRNA-1 | +96 to +115 | ATAGTTGCTGTTGCCAAAAAACATAACCTGTACCGAAGAAGCAGAGATCCAAGCATCTGAgttttagagctagaaatagcaag |
| *tra1* | DHO 1820 | ATG sgRNA-2 | +126 to +107 | ATAGTTGCTGTTGCCAAAAAACATAACCTGTACCGAAGAATGATACATCCATCAGATGCTgttttagagctagaaatagcaag |
| *tra2* | DHO 1593 | 3' loxP insertion / pUG6 | +10950 to +11029 | CTTTGGTGGATTTAGTATCTCAAGCAACTAATCCTCAGCAGCTGGCGCAAATGGATCAACTTTGGCAAGCTTGGCTTTAAAGCTGAAGCTTCGTACGCTGC |
| *tra2* | DHO 1594 | 3' loxP insertion / pUG6 | +11110 to +11030 | ATTTTCTCCTAATAATAAATAATCTGAGCATTCGTAGCAATTCAATATTAGATGAAACCTCAAAACTGATTTGCAATTTGTATAGGGAGACCGGCAGATCC |
| *tra2* | DHO 1615 | 5' loxP insertion / pFA6a | -231 to -166 | TTGTATTTGATAAAAATTACACTCACGTCGATTTCAAATAGCTTCAGTGGCTTCTTGAATAGTGCAATAACTTCGTATAATGTATGCTATACGAAGTTATCGGATCCCCGGGTTAATTAA |
| *tra2* | DHO 1616 | 5' loxP insertion / pFA6a | -86 to -165 | AAAAGCACAGAACGATTACTGTCAGATGTTGTTTAGTTGTTGCTATGTTCAAGGTGAATCGTATAGTGAGGACGCAAAACGAATTCGAGCTCGTTTAAAC |
| *tra2* | DHO 388 | RT-qPCR | +9307 to +9326 | CTCGTTTTCAAGCGTTAGCC |
| *tra2* | DHO 389 | RT-qPCR | +9547 to +9528 | AAGAGCGACAATGAGCCTGT |
| *trk2* | DHO 1711 | RT-qPCR | +1642 to +1661 | TCTTGGTTTGGTTGCGTTCC |

| | | | | | |
|---|---|---|---|---|---|
| *trk2* | DHO | 1712 | RT-qPCR | +1789 to +1770 | ATCGCTGTTCAACGAATAGC |
| *tti2* | DHO | 819 | C-terminal tagging / pFA6a | | TAAAATTGAACATAACTGTTGAGCAACTCGGTGATTACAATGTAATATTGCCACTGCTTGAACCCCTGCAAATCCCACTTGGGGGAGGCGGGGGTGGATACCCATACGATGTTCCTGA |
| *tti2* | DHO | 1222 | C-terminal tagging / pFA6a | | TAAAATTGAACATAACTGTTGAGCAACTCGGTGATTACAATGTAATATTGCCACTGCTTGAACCCCTGCAAATCCCACTTGGGGGAGGCGGGGGTGGA |
| *tti2* | DHO | 820 | Deletion & C-terminal tagging / pFA6a | | TTTTTCGAATAACCATTGTTACCAATTTATCTAGGTTGAAGGTTGGTTAGTAATGAGGAGAGTCTTTCCTTATATGTACCGAATTCGAGCTCGTTTAAAC |
| *tti2* | DHO | 1522 | 3' loxP insertion / pUG75 | +1576 to +1497 | TTTTTCGAATAACCATTGTTACCAATTTATCTAGGTTGAAGGTTGGTTAGTAATGAGGAGAGTCTTTCCTTATATGTACCAGGGAACAAAAGCTTGCATG |
| *tti2* | DHO | 1523 | 3' loxP insertion / pUG75 | +1414 to +1493 | TAAAATTGAACATAACTGTTGAGCAACTCGGTGATTACAATGTAATATTGCCACTGCTTGAACCCCTGCAAATCCCACTTGGTGGATCTGGTGGATCT |
| *tti2* | DHO | 1524 | 5' loxP insertion / pUG6 | -387 to -308 | CTTGGTAATAACTATAAAAAACAATAATGCTTATATACTCAATAGAACTTATATAGCTAAAGCGATAAGGTAAAGTACCTGGTCGACAACCCTTAATATA |
| *tti2* | DHO | 1525 | 5' loxP insertion / pUG6 | -228 to -307 | AATTTAATAAATTAACAATTTACTAGTTTTTAAAGGTATAAAAAGCAAAATTTTTAGTAACTAACATCAATTCATTACACATTAAGGGTTCTCGAGAGCT |
| *vid21* | DHO | 1821 | C-terminal tagging / pFA6a | | GTTCTTTAAAACTAACACCTGAACAAATTCATCAGTTGCAGCAAAGGAAGCAAACTGTACCTACTACTGAAAGGACACAGCGGATCCCCGGGTTAATTAA |
| *vid21* | DHO | 1822 | Deletion & C-terminal tagging / pFA6a | | CTCGCCTCGTAATTTCCTTCTAATATACAACAGCACTCAAAAAAAATACAATAAATCGAAAAATTGCATTGAGGCGATACGAATTCGAGCTCGTTTAAAC |

[a] pFA6a, pKSura4 and DHB plasmids are defined in Materials and Methods

[b] Coordinates are relative to the ATG of each ORF (A defined as +1)
